# Comparative transcriptomic profiling of human conjunctival epithelial cells and macrophages in response to *Chlamydia trachomatis* genovars A and B in early- and mid-infection cycles

**DOI:** 10.1101/2025.11.24.689971

**Authors:** Ehsan Ghasemian, Martin J. Holland

## Abstract

*Chlamydia trachomatis* (Ct), an obligate intracellular bacterium, is the primary infectious cause of blindness through trachoma. Ct undergoes a unique biphasic developmental cycle between infectious elementary bodies and replicating reticulate bodies, manipulating host cells via secreted effector proteins. Whilst previous studies have characterised host-pathogen interactions including transcriptomes of urogenital Ct genovars E and L2, limited studies exist on ocular Ct genovars. This study examined transcriptomic responses of human conjunctival epithelial (HCjE) cells and PMA-differentiated THP-1 macrophages to infection with ocular Ct strains A/2497 or B/Tunis864 (live or heat-inactivated) at 4 and 24 hours post-infection (hpi). Transcriptomic profiling was performed using Lexogen QuantSeq 3’ mRNA-Seq, with differential gene expression analysis conducted using DESeq2. Gene Ontology Biological Process and KEGG pathway enrichment analyses were performed using Gene Set Enrichment Analysis via clusterProfiler to identify strain-specific and cell type-specific transcriptional signatures. HCjE cells exhibited progressive transcriptional activation, with differentially expressed genes (DEGs) increasing from 4 to 24 hpi (144 to 259; *P* = 0.0458), whilst THP-1 macrophages showed temporal attenuation (391 to 154; *P* < 0.0001). B/Tunis864 consistently elicited higher responses than A/2497 in HCjE cells at both time points (4 hpi: 152 *vs*. 54, *P* = 0.0003; 24 hpi: 259 *vs*. 83, *P* < 0.0001). Conversely, THP-1 macrophages showed higher responses to A/2497 than B/Tunis864 at both time points (4 hpi: 599 *vs*. 376, *P* < 0.0001; 24 hpi: 166 *vs*. 114, *P* = 0.0221). HCjE cells demonstrated markedly higher proportions of strain-specific DEGs and pathways compared to macrophages. B/Tunis864 infection in HCjE cells induced pronounced interferon-stimulated gene signatures, particularly at 24 hpi. This study revealed contrasting temporal patterns: THP-1 macrophages showed peak-then-decline responses, whilst HCjE cells exhibited progressive activation to 24 hpi. HCjE cells demonstrated predominantly strain-specific responses, whereas macrophages deployed strain-invariant programmes. B/Tunis864’s enhanced interferon-stimulated gene and inflammatory pathway activation in HCjE cells may suggest molecular hints for genovar B-associated trachoma severity.

## Introduction

*Chlamydia trachomatis* (Ct) is an obligate intracellular bacterial pathogen that primarily infects ocular and urogenital mucosal epithelia [1,2]. Ct is the etiological agent of trachoma, which represents the leading infectious cause of preventable blindness worldwide, and is also responsible for the most prevalent bacterial sexually transmitted infection [3,4]. The pathological sequelae of chlamydial infection result from dysregulated host inflammatory and immune responses that drive chronic inflammation, progressive tissue remodelling, and ultimately fibrotic scarring of the upper reproductive tract and conjunctival tissues [1,5,6]. These pathological processes are mediated through host-pathogen interactions involving multiple mechanisms. Cell-autonomous immune responses, such as IL-1R-dependent pathways, play a dual role: whilst promoting bacterial clearance, they simultaneously inflict collateral tissue damage that contributes to pathology [7]. Additionally, Ct infection may induce epithelial-mesenchymal transition (EMT) and stimulate production of extracellular matrix components and collagen I in host epithelial cells, establishing a pro-fibrotic programme that operates concurrently with inflammatory responses [8–10]. This dual activation of pro-inflammatory and pro-fibrotic signalling pathways in infected epithelial cells represents an expanded cellular paradigm of chlamydial pathogenesis, wherein tissue scarring results not only from collateral inflammatory damage but also from direct pathogen-induced fibrotic remodelling [10]. Furthermore, Ct actively manipulates host immune signalling through secreted bacterial effector proteins, exemplified by the deubiquitinating protease ChlaDub1, which inhibits NF-κB activation and disrupts normal immune cell responses, thereby facilitating bacterial survival and persistence [11–13].

Ct has a unique biphasic developmental cycle, alternating between two morphologically and functionally distinct forms: metabolically inactive, infectious elementary bodies (EBs) and metabolically active, replicating reticulate bodies (RBs) [14]. The chlamydial developmental cycle can be temporally divided into early (1-8 hours post-infection (hpi)), mid (8-48 hpi), and late (48-72 hpi) phases based on differential gene expression patterns, replicative activity, and host-pathogen interactions [15–18]. Infection is initiated when EBs attach to and invade susceptible host cells through receptor-mediated endocytosis [15,19]. Following internalisation, EBs establish residence within membrane-bound inclusion vacuoles that actively evade phagolysosomal fusion, creating a protective intracellular niche where they differentiate into metabolically active RBs. After multiple rounds of binary fission, RBs asynchronously re-differentiate back into infectious EBs, which are subsequently released through either inclusion extrusion or host cell lysis to reinitiate the infection cycle [15,19]. Throughout this developmental process, Ct employs its type III secretion system (T3SS) to translocate effector proteins directly from the bacterial cytoplasm into the host cell cytosol [20], and secrete virulence factors that manipulate multiple aspects of host cell biology: TarP and TmeA coordinate actin cytoskeleton reorganisation to facilitate invasion; IncD and IncV establish ER-inclusion membrane contact sites essential for intracellular survival and replication; whilst GarD, CpoS, and chlamydial protease-like activity factor (CPAF) collectively subvert both innate and adaptive immune responses to ensure bacterial persistence [15,19,21,22].

Previous studies by Hayward *et al*. [23,24] provided insights into host-pathogen interactions during Ct genovar E (CtE) infection through the application of single-cell RNA sequencing and dual RNA-seq approaches to simultaneously analyse host and bacterial transcriptomes. These studies demonstrated that adherence and uptake of Ct rapidly triggers transcriptional reprogramming in infected host cells [23,24]. This work revealed the role of Ct in subverting host immune responses through multiple mechanisms, including downregulation of antimicrobial peptides and mucin expression, attenuation of innate immune signalling and modulation of pathways controlling immune cell recruitment and activation [25]. It was hypothesised that these transcriptional alterations may contribute to progressive tissue damage and pathological sequelae [25]. Furthermore, *in vitro* studies utilising genital CtE and genovar L2 have elucidated the temporal dynamics of host transcriptional modulation, revealing differential regulation of genes involved in metallothionein function, cell cycle control, innate immunity, cytoskeletal organisation, lipid biosynthesis, and cellular stress responses compared to mock-infected controls [23,26,27].

Ct genovars A (CtA), B (CtB) and C are the causative agents of blinding trachoma [3,28], and may exhibit different molecular strategies for epithelial cell interaction and immune evasion [14,29–32]. Previous investigations have documented distinct bacterial load kinetics, disease severity, infection duration, and genomic evolutionary pressures between CtA and CtB [33–42]. Despite advances in understanding Ct-host interactions, our knowledge of how these ocular genovars modulate host cell transcriptomes remains limited. In this study, we conducted transcriptomic analysis of phorbol myristate acetate (PMA)-activated THP-1 cells and immortalised human conjunctival epithelial (HCjE) cells infected with either live or heat-inactivated (HIA) Ct strain A/2497 or B/Tunis864 at 4 and 24 hpi (Fig. 1). HCjE cells, are immortalised conjunctival epithelial cells and represent the primary target tissue for ocular Ct infection [27,43,44], whilst PMA-activated THP-1 cells model the monocyte-derived macrophages that are recruited to the site of infection [45–47]. It is known that the initial engagement between macrophages and Ct may determine the overall outcome of the infection since if intracellular elimination in macrophages fails, macrophages may be used as Trojan horses for dissemination of Ct [45,48]. Through this study we aim to: (*i*) evaluate the responsiveness of HCjE and THP-1 cells to Ct strains A/2497 and B/Tunis864 infection, representing CtA and CtB, respectively, (*ii*) identify differential host cellular pathway enrichment patterns in response to A/2497 *vs*. B/Tunis864 infection, (*iii*) determine transcriptional specifications associated with early- and mid-developmental stages of Ct, and (*iv*) suggesting genes and pathways underlying the pathogenesis and varied clinical outcomes of trachoma in relation to Ct genovar.

**Fig. 1.**
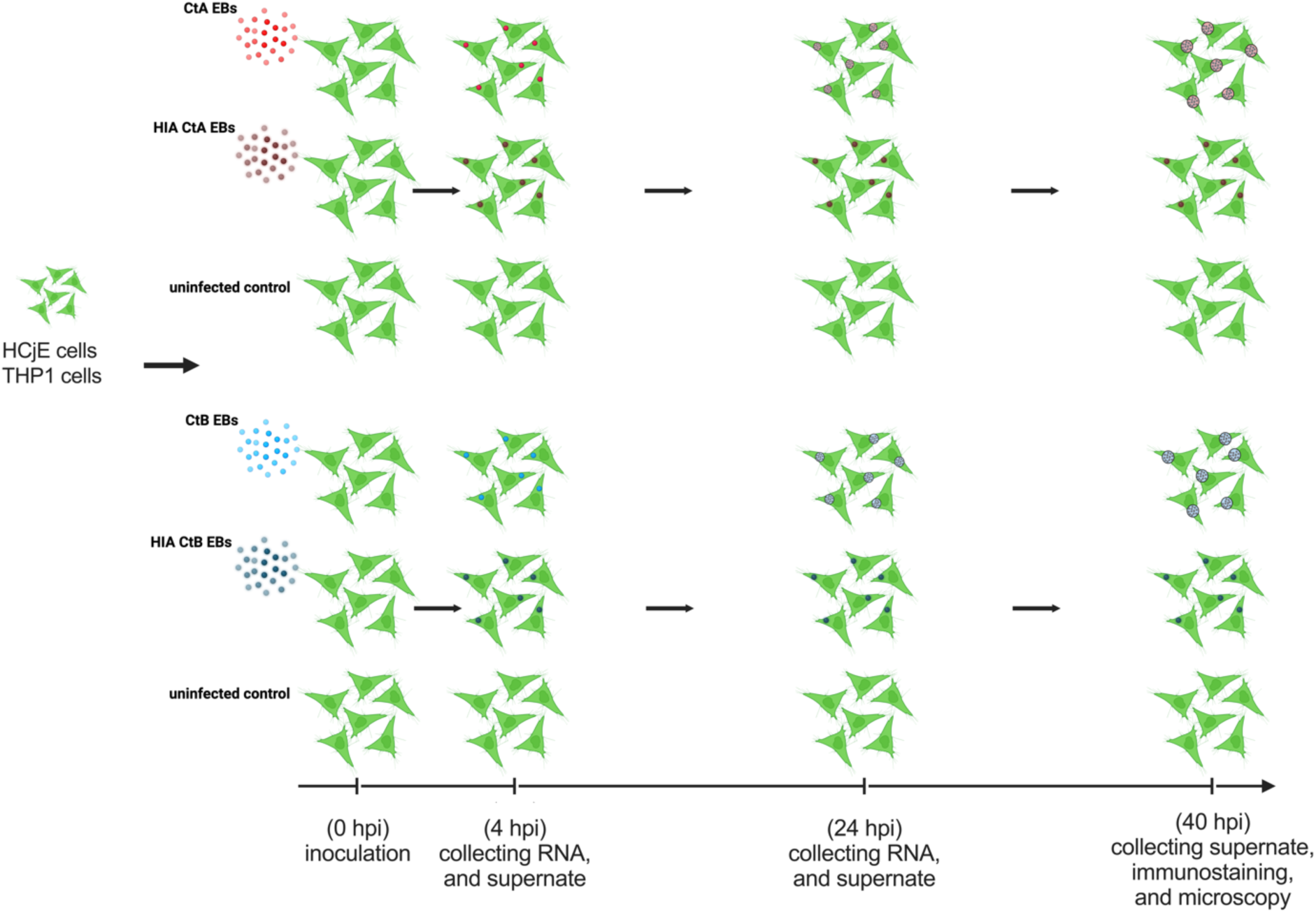
Experimental Design. HCjE and THP-1 cells were inoculated with live or HIA CtA or CtB, with uninfected cells serving as negative controls. RNA was extracted at 4 and 24 hpi, and chlamydial infection was assessed by microscopy at 40 hpi. Color coding: live CtA (red), HIA CtA (dark red), live CtB (blue), HIA CtB (dark blue).

## Materials and methods

### Bacterial stock preparation

HeLa cells monolayers, kindly provided by Professor David Allen (London School of Hygiene and Tropical Medicine), were infected with Ct strains A/2497 [49] and B/Tunis864 [50]. For bacterial stock preparation, EBs were propagated in HeLa cell monolayers and subsequently purified using Gastrografin density gradient centrifugation [51]. Purified Ct EB stocks were titrated by serial dilution and reinfection of HeLa cell monolayers, with quantification performed using Pathfinder anti-chlamydial lipopolysaccharide (LPS) immunofluorescent staining (Bio-Rad). Fluorescent imaging and counting of inclusion-forming units (IFUs) were conducted using a Nikon ECLIPSE Ti microscope (Nikon Instruments Inc.). The purified Ct EBs were resuspended in sucrose phosphate glutamate (SPG) buffer and stored at −70°C until use.

### Inoculation of HCjE and THP-1 cells

HCjE cells, kindly provided by Professor Ilene K. Gipson (Harvard Medical School), were cultured as monolayers in 6-well plates until reaching 90% confluency using Keratinocyte Serum-Free Medium (SFM) (Gibco) supplemented with 25 ug/mL Bovine Pituitary Extract (BPE), 0.2 ng/mL epidermal growth factor (EGF), 0.4 mM CaCl_2_ (Merck), 25 ug/mL gentamicin gentamicin (Gibco), 20 ug/mL vancomycin (Fisher scientific), and 50 ug/mL amphotericin B (Gibco). THP-1 cells (a human monocyte cell line), kindly provided by a colleague at the London School of Hygiene and Tropical Medicine, were maintained in RPMI 1640 medium (Gibco) supplemented with 10% fetal bovine serum (FBS) (Gibco), 2 mM L-Glutamine (Gibco), 0.05 mM 2-mercaptoethanol (Merck), 25 ug/mL gentamicin (Gibco), 20 ug/mL vancomycin (Fisher scientific), and 50 ug/mL amphotericin B (Gibco). For macrophage differentiation, THP-1 cells were seeded in 6-well plates at 1x 10^6^ cells per well and treated with 0.16 μg/mL phorbol 12-myristate 13-acetate (PMA) (Merck) for 72 h to induce differentiation into macrophage-like cells.

Inoculation medium was prepared specifically for each cell type. For HCjE cells, Dulbecco’s Modified Eagle Medium (DMEM)/F-12 (Gibco) was supplemented with 1% gentamicin (Gibco) and 1% L-glutamine (Gibco). For THP-1 cells, RPMI-1640 (Gibco) medium was supplemented with 10% FBS (Gibco), 2 mM L-glutamine (Gibco), and 1% gentamicin (Gibco). Prior to infection, cell monolayers were washed three times with Hank’s Balanced Salt Solution (HBSS) (Merck). Each cell line was inoculated in biological triplicate with: (*i*) live Ct at MOI 5, (*ii*) HIA Ct at MOI 5, or (*iii*) inoculation medium alone as a mock-infected control. Infection was synchronised by centrifugation at 400 x g for 30 min, followed by incubation for 2 h at 37°C with 5% CO_2_. At 2 hpi, cells were washed three times with HBSS to remove extracellular EBs, and fresh inoculation medium was added. For transcriptomic analysis, cells were lysed at 4 and 24 hpi using Buffer SKP from the NORGEN BIOTEK DNA/RNA purification kit. Additional samples were collected at 40 hpi and fixed with methanol (Merck) for subsequent immunofluorescent analysis using the Pathfinder anti-chlamydial lipopolysaccharide (LPS) immunofluorescent staining (Fig. 1). Prior to all sample collection timepoints, cells were washed three times with phosphate-buffered saline (Gibco) to remove dead or unattached cells.

### Library preparation and sequencing

Cell lysates collected at 4 and 24 hpi were subjected to RNA extraction using the NORGEN BIOTEK DNA/RNA Purification Kit according to the manufacturer’s protocols for total RNA purification from cellular lysates.

For RNA sequencing, cDNA library preparation was performed using the Lexogen QuantSeq 3’ mRNA-Seq Library Prep Kit for Illumina (FWD) following the manufacturer’s protocol. Double-stranded cDNA libraries were purified using magnetic bead-based purification. To optimise amplification efficiency, the number of PCR cycles for library amplification was determined through quantitative PCR using the Lexogen PCR Add-on Kit for Illumina. Sample-specific barcoding was achieved using the Lexogen UDI 12 nt Unique Dual Indexing Add-on Kit to incorporate unique dual indices. Following final magnetic bead purification, libraries were quantified and pooled in equimolar ratios to ensure balanced representation during multiplexed sequencing. Pooled libraries were submitted to Lexogen for single-end sequencing on the NextSeq 2000 platform (Illumina).

### Data processing and differential expression analysis

Raw single-end sequencing reads were processed through a standardised bioinformatics pipeline. Adapter trimming and quality filtering were performed using Cutadapt (version 1.18), followed by quality assessment using FastQC (v0.11.7). Trimmed reads were aligned to the human reference genome (GRCh38, Ensembl release 107) using STAR aligner (v2.6.1a), and gene-level read quantification was performed using featureCounts (v1.6.4). Alignment quality metrics, including the percentage of uniquely mapped reads, were compiled using MultiQC (v1.5).

Differential gene expression analysis was conducted in R (v3.6.0) using RStudio (v2024.04.02) with the tidyverse package (v1.2.1) for data manipulation and visualisation. To ensure robust statistical analysis, genes with insufficient read coverage (present in fewer than 3 samples with at least 10 counts) were filtered prior to analysis. Differential expression analysis was performed using the DESeq2 package (v1.18.1) following the standard workflow, with count data normalisation performed using DESeq2’s median-of-ratios method, which calculates sample-specific size factors by taking the median ratio of each sample’s counts to the geometric mean across all samples for each gene to account for differences in sequencing depth and RNA composition. Genes were classified as differentially expressed if they met the criteria of log2 Fold Change (log2FC) > 1 and adjusted *P*-value < 0.05 (Benjamini-Hochberg multiple testing correction). Ensembl gene identifiers were converted to HGNC gene symbols using the org.Hs.eg.db annotation package (v3.8.2). Statistical significance testing for differential gene expression counts between treatment conditions was performed using Fisher’s exact test.

### Data visualisation and exploratory analysis

Principal component analysis (PCA) was performed to visualise sample clustering and assess global transcriptional patterns across experimental conditions. Normalised count data for differentially expressed genes were subjected to z-score scaling (mean-centred and scaled to unit variance) to standardise gene expression values across samples prior to dimensionality reduction. PCA was prepared using the prcomp() function in R. Sample clustering was visualised using ggplot2 (v3.3.2), and confidence ellipses drawn around each group using the ggforce package (v0.3.2) to highlight cluster boundaries. Sample labels were added using the ggrepel (v0.9.6), and a custom colour palette was generated using RColorBrewer (v1.1.3).

Gene expression heatmaps were generated to visualise hierarchical clustering patterns and identify co-expressed gene modules across experimental conditions using the pheatmap package (v1.0.12). Normalised count data were partitioned into experimental groups and subsequently merged and subjected to z-score scaling (mean-centred and scaled to unit variance) to standardise expression values and enable comparison across genes with different baseline expression levels. Hierarchical clustering was performed using Euclidean distance metrics. Dendrograms were cut to form 4 distinct clusters for both genes and samples to identify major expression patterns, with a custom colour palette.

### Pathway enrichment analysis

Functional enrichment analysis was performed to identify biological pathways and processes associated with differentially expressed genes using Gene Set Enrichment Analysis (GSEA) with the clusterProfiler (v4.12.6), fgsea (v1.30.0), and org.Hs.eg.db packages in R. For Gene Ontology (GO) Biological Process (BP) analysis, genes were ranked based on log2FC values to preserve the magnitude and direction of expression changes across the entire transcriptome. GO BP and associated gene sets were retrieved from the org.Hs.eg.db annotation database using the AnnotationDbi::select() function from the AnnotationDbi package (v1.66.0), with analysis restricted to gene sets containing 15-1000 genes to ensure statistical power whilst excluding overly broad or highly specific categories. GSEA was conducted using the gseGO() function from clusterProfiler (v4.12.6), focussing on BP ontology terms with a significance threshold of adjusted *P*-value < 0.05 using Benjamini-Hochberg multiple testing correction. To reduce redundancy amongst significantly enriched GO BP, pathway consolidation was performed using the collapsePathways function from fgsea, which retains the most representative pathways from clusters of overlapping gene sets based on pathway similarity.

For KEGG pathway analysis, HGNC gene symbols were converted to ENTREZ identifiers using the bitr() function from clusterProfiler, with genes lacking valid ENTREZ IDs or duplicate mappings removed to maintain analytical integrity. Pre-ranked gene lists were analysed using the gseKEGG() function against the human KEGG database (organism = “hsa”) with identical gene set size restrictions (15-1000 genes). For KEGG analyses, pathways with adjusted *P*-values < 0.05 (Benjamini-Hochberg multiple testing correction) were considered statistically significant. Pathways were classified as upregulated or downregulated based on their normalised enrichment scores.

### Pathway visualisation and comparative analysis

Enriched GO BP and KEGG pathway data were visualised using customised combined bar and dot plots generated with ggplot2. Gene ratios and adjusted *P*-values were processed to create bar plots displaying enrichment scores. Network plots were generated using the cnetplot() function from clusterProfiler to visualise relationships between significantly enriched pathways and their constituent genes, with colour gradients representing log2FC. Plot customisation was achieved using ggplot2 and gridExtra (v2.3) packages.

To identify pathway enrichment patterns across treatment conditions, intersection analysis was performed using Venn diagrams generated with the ggvenn (v0.1.10) package. Pathways were systematically categorised into four functional groups based on inoculation conditions: (*i*) CtA-specific pathways (unique to live CtA or shared between live and HIA CtA), (*ii*) CtB-specific pathways (unique to live CtB or shared between live and HIA CtB), (*iii*) core pathways (shared amongst live CtA and live CtB; live CtA and HIA CtB; live CtA, and HIA CtA and live CtB; live CtB and HIA CtA; live CtB, HIA CtB and live CtA) and (*iv*) HIA pathways (specific to HIA CtA, HIA CtB, or shared between both HIA CtA and HIA CtB). Statistical significance of pathway distribution across inoculation conditions was assessed using Poisson distribution testing.

### Pathway-associated gene expression analysis

Genes associated with strain-specific significantly enriched GO BP and KEGG pathways were identified and subjected to differential expression analysis. Differential expression analysis was performed on DESeq2 (v1.44.0) data. For comparative analysis between strains, genes were filtered based on statistical significance (adjusted *P* < 0.05, Benjamini-Hochberg correction) in at least one strain. Expression patterns were then characterised based on Log2FC differences between strains using a 1.5-fold threshold (Log2FC difference = 0.585).

Genes were categorised into six response patterns: (*i*) CtA-specific - significant only in CtA (adjusted *P* < 0.05); (*ii*) CtB-specific - significant only in CtB (adjusted *P* < 0.05); (*iii*) Concordant - significant in at least one strain with the same regulatory direction and Log2FC difference < 0.585; (*iv*) CtA-dominant - significant in at least one strain with the same regulatory direction and CtA exhibiting ≥1.5-fold greater magnitude change; (*v*) CtB-dominant - significant in at least one strain with the same regulatory direction and CtB exhibiting ≥1.5-fold greater magnitude change; (*vi*) Opposite - significant in at least one strain with opposing regulatory directions (both strains showing absolute Log2FC > 0.585). Effect sizes were calculated as Cohen’s d using the approximation d = Log2FC/lfcSE, where lfcSE represents the standard error of the Log2FC from DESeq2 output. Mean effect size for comparative analysis was calculated as the average of absolute Cohen’s d values: (Cohen’s d_CtA + Cohen’s d_CtB)/2.

Data visualisation included hierarchical clustering heatmaps generated using pheatmap (v1.0.12) with correlation-based distance metrics and Ward.D2 clustering method. Two heatmaps were generated: (*i*) all pathway-associated genes to provide comprehensive overview, and (*ii*) only significantly differentially expressed genes (adjusted *P* < 0.05 in CtA, CtB, or both) to focus on strain-specific responses. Heatmaps displayed mean normalised expression values across four conditions: CtA control, CtA infected, CtB control, and CtB infected, with row annotations indicating pathway origin and significance levels.

## Results

### Ct infection and RNA-seq data quality assessment

HCjE and THP-1 cells were infected with Ct strains A/2497 and B/Tunis864, and confirmed by microscopy (Fig. S1, Fig. S2). RNA-seq data quality metrics before and after trimming are summarised in Tables S1 and S2, and Figure S3. Following quality trimming and alignment using STAR, we obtained an average of 2,809,517 ± 812,089 (mean ± SD) uniquely mapped reads per sample, corresponding to 11,674 ± 1,909 detected genes. Detailed alignment statistics, including the number and percentage of uniquely mapped reads, are provided in Table S3 and Figure S4.

### Transcriptomic responses in HCjE and THP-1 cells following Ct infection

PCA of all experimental conditions revealed that PC1 and PC2 explained 28.6% and 9.4% of the total variance, respectively, primarily capturing variation associated with (*i*) cell type and (*ii*) time post-inoculation (Fig. 2a). Hierarchical clustering analysis, based on normalised mean counts for each condition, supported these findings, resulting in clustering by cell line and inoculation time point (Fig. 2d). Cell line-specific PCA showed that PC1 and PC2 accounted for 22.8% and 14.4% of variance in HCjE cells, and 27.8% and 11.3% in THP-1 cells, respectively. These components appeared to reflect differences in, inoculation time point and inoculation condition (Fig. 2b, Fig. 2c).

**Fig. 2.**
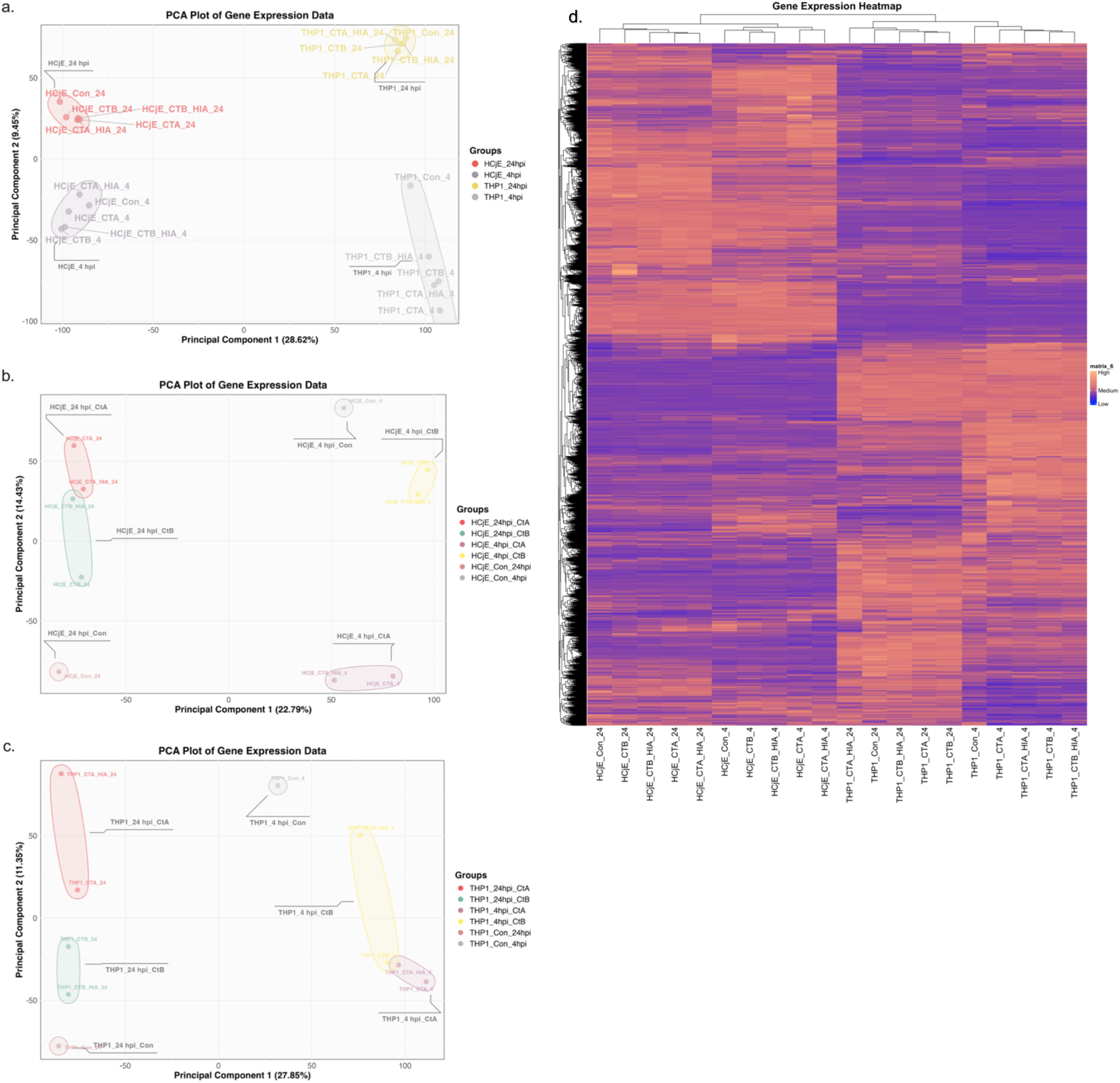
PCA and hierarchical clustering of gene expression data. (**a**) PCA plot showing sample separation by cell type (HCjE *vs*. THP-1) and time point (4 *vs*. 24 hpi) along PC1 and PC2. (**b**) PCA plot for HCjE cells demonstrating separation by time point and inoculation condition. (**c**) PCA plot for THP-1 cells showing similar separation by time point and inoculation condition. (**d**) Heatmap of normalised expression levels for 12,701 genes across all conditions. Data were normalised to the mean expression across samples, with hierarchical clustering applied to both genes (rows) and samples (columns) using complete linkage and Euclidean distance.

In HCjE cells, DEG numbers were significantly higher at 4 hpi compared to 24 hpi (mean DEGs: 144 *vs*. 100; *P* = 0.0458) (Fig. S5a). THP-1 cells showed a significant reduction in the number of DEGs from 4 hpi to 24 hpi (mean DEGs: 154 *vs*. 391; *P* < 0.0001) (Fig. S5b). In HCjE cells infected with A/2497 there were significantly fewer DEGs than B/Tunis864 at both time points (4 hpi: 54 *vs*. 152, *P* = 0.0003; 24 hpi: 83 *vs*. 259, *P* < 0.0001) (Fig. S5a). Conversely, in THP-1 cells, A/2497 infection resulted in significantly more DEGs than B/Tunis864 at both time points (4 hpi: 599 *vs*. 376, *P* < 0.0001; 24 hpi: 166 *vs*. 114, *P* = 0.0221) (Fig. S5b).

Analysis of DEG intersections across the eight inoculation conditions within each cell line revealed distinct patterns (Fig. S5). Of the total DEGs identified (HCjE: 414 at 4 hpi, 329 at 24 hpi; THP-1: 794 at 4 hpi, 369 at 24 hpi), THP-1 cells showed a significantly higher proportion of overlapping genes at each time point (4 hpi: 50.1%; 24 hpi: 36.9%) compared to HCjE cells (4 hpi: 24.6%; 24 hpi: 14%) (both *P* < 0.0001) (Fig. S5). The highest numbers of unique DEGs were observed in HCjE cells infected with live B/Tunis864 at 24 hpi (29.1%) and THP-1 cells infected with live A/2497 at 4 hpi (20.7%) (Fig. S5).

### Pathway enrichment analysis in HCjE and THP-1 cells

There was a significant increase in the mean number of enriched pathways from 4 hpi to 24 hpi for both GO BP (2.5 to 13.5; *P* = 0.003) and KEGG (1.75 to 14.75; *P* < 0.0001) in HCjE cells (Fig. 3). At 4 hpi, 10 GO BP and 7 KEGG pathways were significantly enriched, with no pathways shared across all inoculation conditions (Fig. 4a, Fig. 4b). By 24 hpi, pathway enrichment had increased to 54 GO BP and 59 KEGG pathways (Fig. 3) out of which no GO BP and 7 KEGG pathways (11.9%) were conserved across all inoculation conditions (Fig. 4c, Fig. 4d). Live B/Tunis864 infection resulted in significantly more enriched GO BP pathways than live A/2497 infection (*P* < 0.0001) (Fig. 3).

**Fig. 3.**
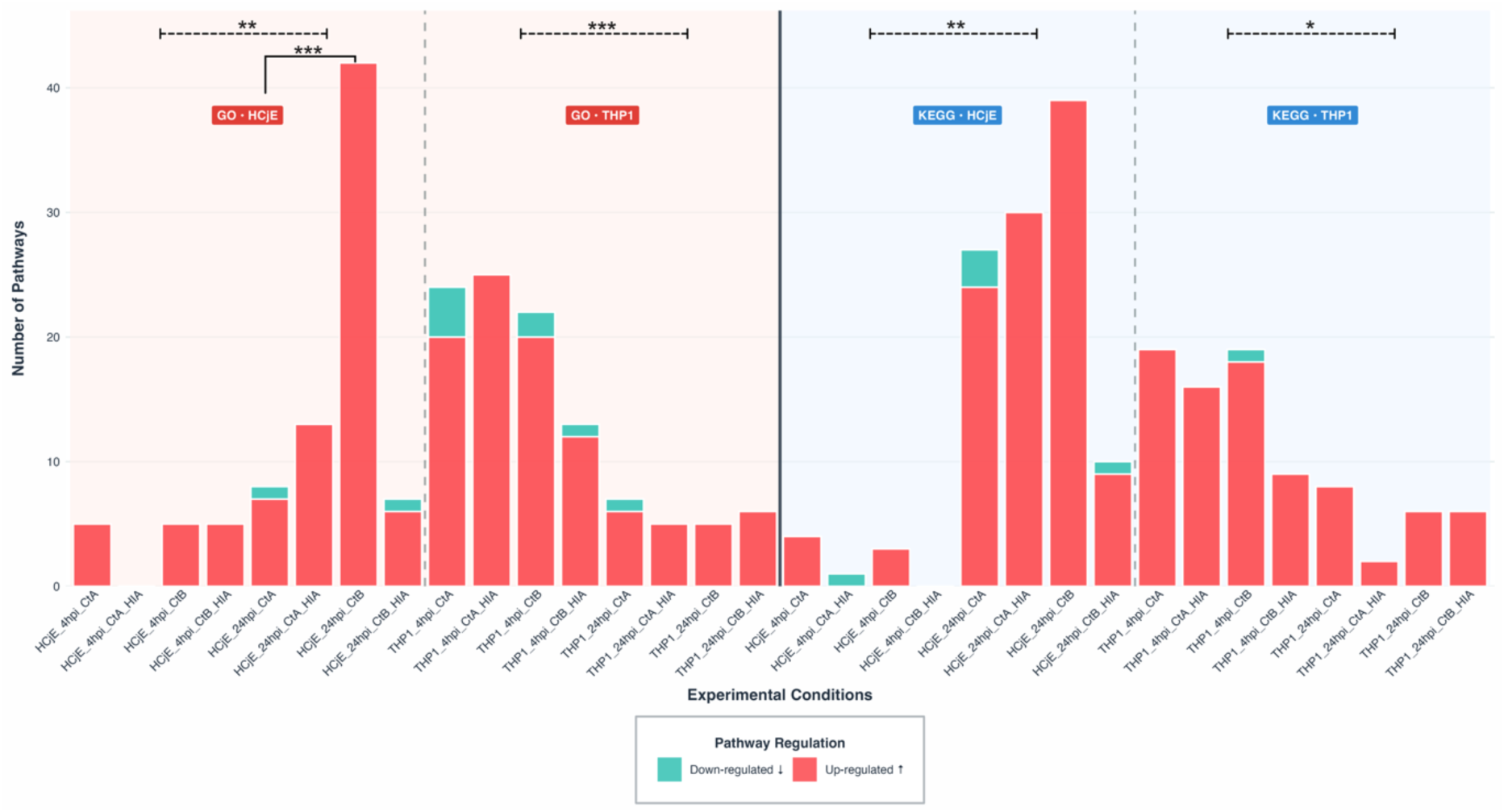
Bar plots showing the number of enriched GO BP (a) and KEGG (b) pathways. The bars represent the total number of enriched pathways in HCjE and THP-1 cells inoculated with CtA and CtB, either live or HIA. Upregulated pathways are shown in red, whilst downregulated pathways are shown in blue. Vertical dashed lines separate conditions associated with HCjE and THP-1 cells, whereas horizontal dashed lines group conditions based on inoculation time points. Statistical significance levels are indicated by asterisks (* *p* < 0.05; ** *p* < 0.01; *** *p* < 0.001).

**Fig. 4.**
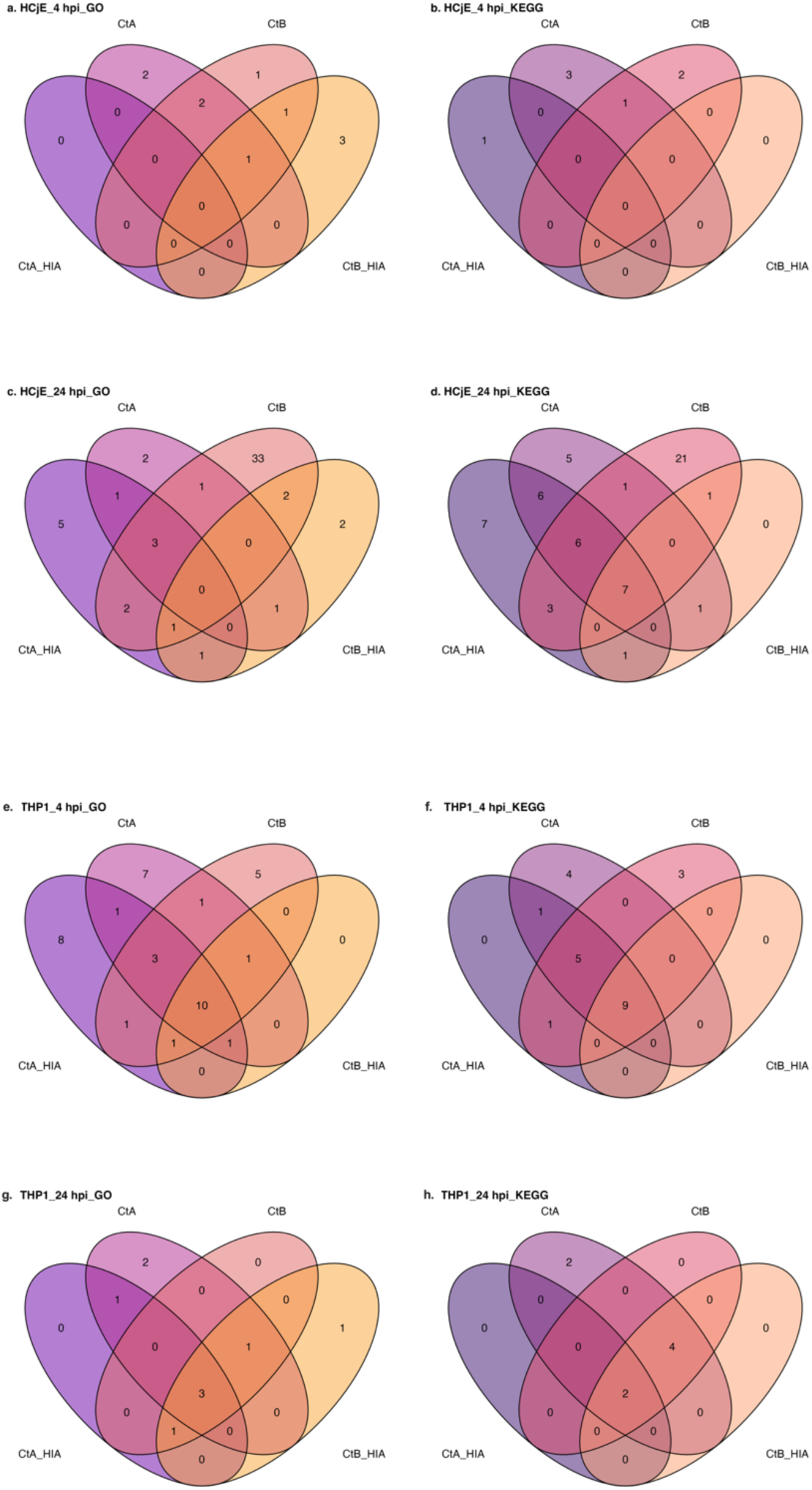
Venn diagrams showing pathway overlap between Ct inoculation conditions in HCjE and THP-1 cells. Diagrams compare GO BP and KEGG pathways identified in response to live CtA, HIA CtA, live CtB, and HIA CtB at 4 and 24 hpi. HCjE cells: GO BP pathways at 4 hpi (**a**), KEGG pathways at 4 hpi (**b**), GO BP pathways at 24 hpi (**c**), KEGG pathways at 24 hpi (**d**). THP-1 cells: GO BP pathways at 4 hpi (**e**), KEGG pathways at 4 hpi (**f**), GO BP pathways at 24 hpi (**g**), KEGG pathways at 24 hpi (**h**).

In contrast to HCjE cells, THP-1 cells exhibited a significant decrease in pathway enrichment from 4 hpi to 24 hpi for both GO BP (9.75 to 2.25; *P* = 0.006) and KEGG (5.75 to 2; *P* = 0.04) analyses (Fig. 3). At 4 hpi, 39 GO BP and 23 KEGG pathways were enriched, with overlap across conditions: 10 GO BP pathways (25.6%) and 9 KEGG pathways (39.1%) were shared across all conditions (Fig. 4e, Fig. 4f). By 24 hpi, pathway enrichment had reduced to 9 GO BP and 8 KEGG pathways, with 3 GO BP pathways (33.3%) and 2 KEGG pathways (25%) conserved across all conditions (Fig. 4g, Fig. 4h).

### Pathway responses in HCjE cells

At 4 hpi in HCjE cells, pathway enrichment analysis revealed distinct strain-specific responses alongside shared core pathways (Fig. 4a, Fig. 4b). Both strains upregulated core pathways involved in stress signalling (GO:0140467), metabolic reprogramming (GO:0045444), inflammation (GO:0006954), and transcriptional dysregulation (hsa05202). B/Tunis864 consistently exhibited higher enrichment scores across these pathways compared to A/2497 (Table 1-1, Fig. S6a-d).

**Table 1-1.** Pathways identified by Gene Set Enrichment Analysis (GO BP and KEGG) for HCjE cells inoculated with Ct strains A/2497 and B/Tunis864 at 4 hpi.

| ID | Description | enrichment score | enrichment score | enrichment score | enrichment score |
| --- | --- | --- | --- | --- | --- |
| **CtA\*** |  |  |  |  |  |
| GO:0048661 | positive regulation of smooth muscle cell proliferation | 0.54 |  |  |  |
| GO:0045087 | innate immune response | 0.33 |  |  |  |
| hsa04061 | Viral protein interaction with cytokine and cytokine receptor | 0.65 |  |  |  |
| hsa05323 | Rheumatoid arthritis | 0.56 |  |  |  |
| hsa04060 | Cytokine-cytokine receptor interaction | 0.49 |  |  |  |
| **CtB<sup>§</sup>** |  |  |  |  |  |
| GO:0032731 | positive regulation of interleukin-1 beta production | 0.61 |  |  |  |
| GO:0010628 | positive regulation of gene expression | 0.36 |  |  |  |
| hsa05031 | Amphetamine addiction | 0.63 |  |  |  |
| hsa05322 | Systemic lupus erythematosus | 0.63 |  |  |  |
| **Core<sup>#</sup>** |  |  |  |  |  |
|  |  | **CtA** | **CtB** | **CtA_HIA** | **CtB_HIA** |
| GO:0140467 | integrated stress response signaling | 0.62 | 0.64 | NA | NA |
| GO:0045444 | fat cell differentiation | 0.45 | 0.50 | NA | NA |
| GO:0006954 | inflammatory response | 0.39 | 0.43 | NA | NA |
| hsa05202 | Transcriptional misregulation in cancer | 0.476 | 0.488 | NA | NA |
| **HIA<sup>†</sup>** |  |  |  |  |  |
|  |  |  |  | **CtA_HIA** | **CtB_HIA** |
| GO:1902895 | positive regulation of miRNA transcription |  |  | NA | 0.64 |
| GO:0006334 | nucleosome assembly |  |  | NA | 0.59 |
| GO:0000122 | negative regulation of transcription by RNA polymerase II |  |  | NA | 0.38 |
| hsa05416 | Viral myocarditis |  |  | -0.61 | NA |

**Table 1-2.**
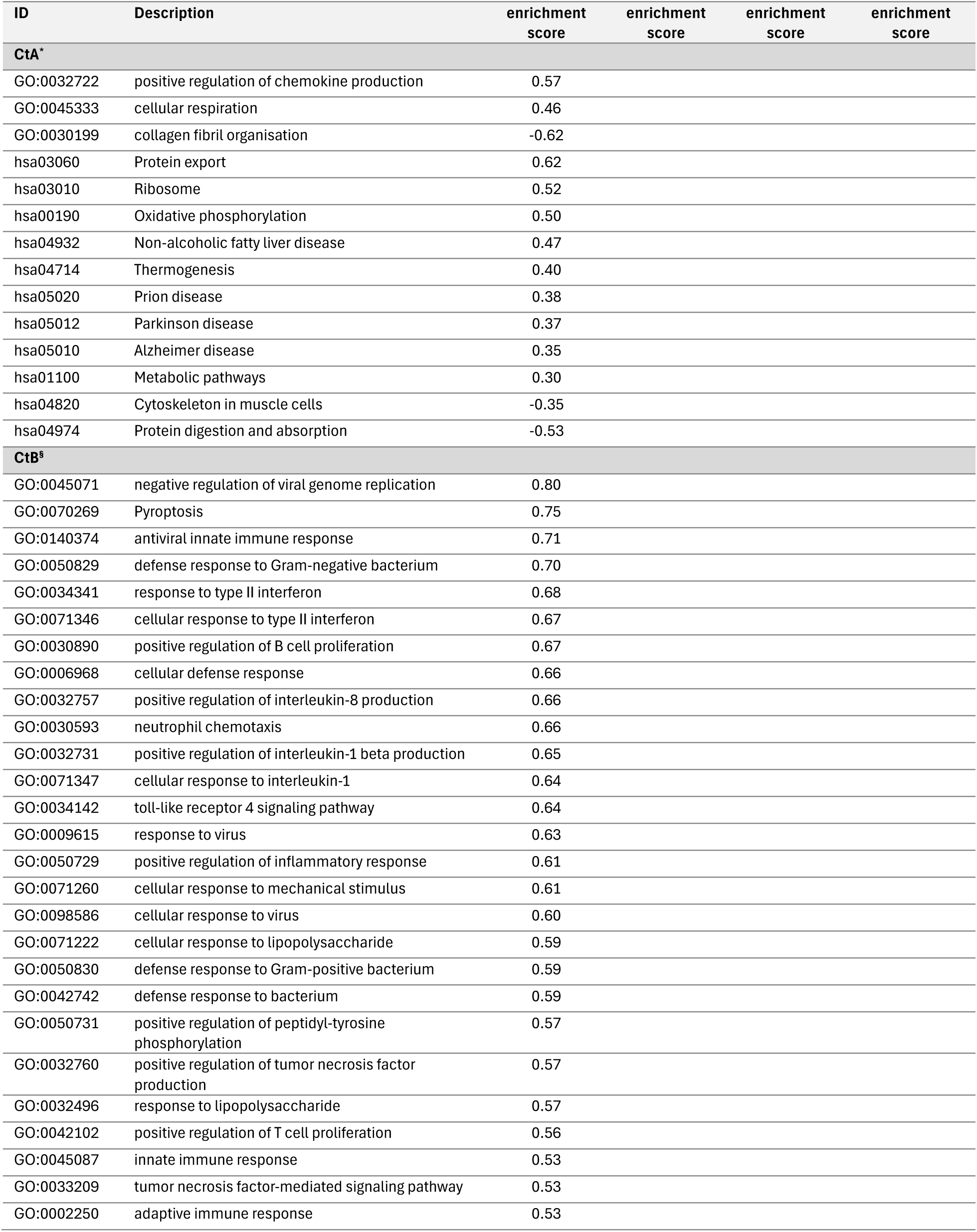
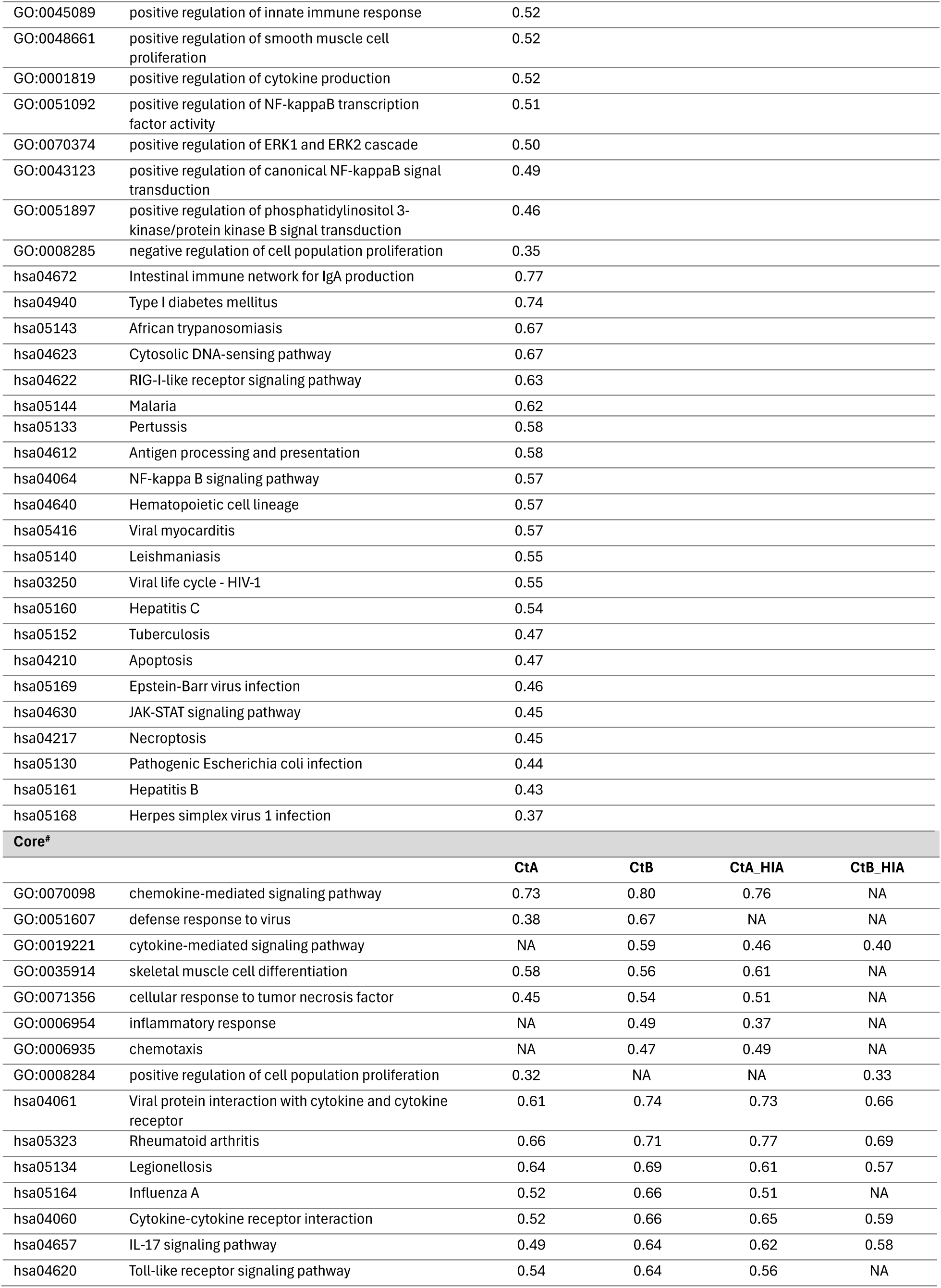
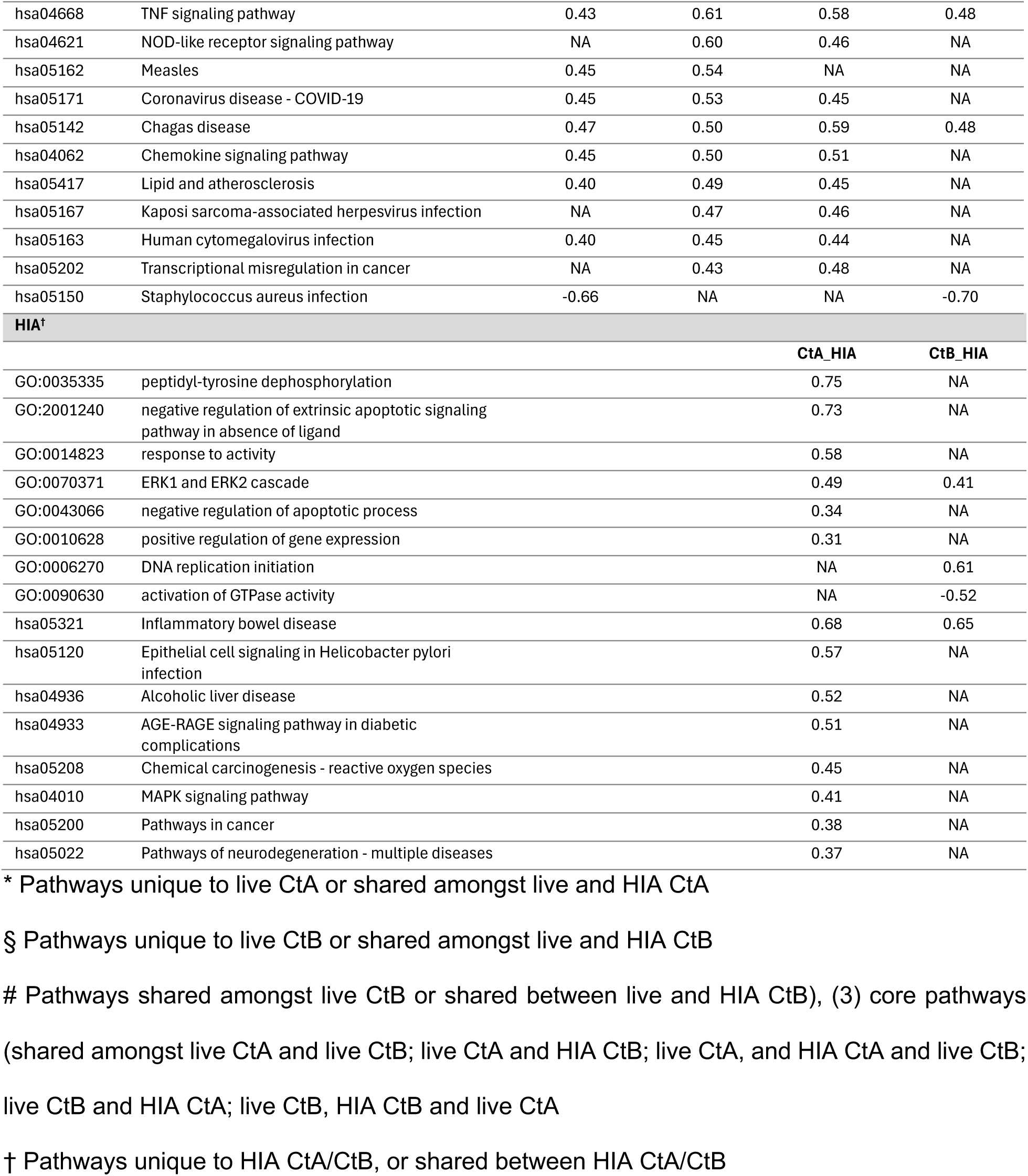
Pathways identified by Gene Set Enrichment Analysis (GO BP and KEGG) for HCjE cells inoculated with Ct strains A/2497 and B/Tunis864 at 24 hpi.

Each strain induced two unique GO BP pathways, whilst, A/2497 uniquely enriched three KEGG pathways and B/Tunis864 enriched two (Table 1-1, Fig. 4a, Fig. 4b). There were functional differences in the pathways activated through each Ct strain: A/2497 activated immune signalling pathways, including innate immune response (GO:0045087), cytokine signalling (hsa04060 and hsa04061), and tissue remodelling (GO:0048661) (Table 1-1, Fig. S6e, Fig. S6g). In contrast, B/Tunis864 induced more targeted inflammatory responses, notably interleukin-1 beta production (GO:0032731), autoimmune-like mechanisms (hsa05322), and neuro-immune interactions (hsa05031) (Table 1-1, Fig. S6f, Fig. S6h).

By 24 hpi, pathway enrichment had diverged markedly between strains. A/2497 uniquely enriched two GO BP pathways compared to 35 for B/Tunis864, whilst eight GO BP pathways remained commonly enriched (Table 1-2, Fig. 4c). For KEGG, A/2497 enriched 11 unique pathways, B/Tunis864 enriched 22, and 18 were shared between strains (Table 1-2, Fig. 4d). The shared core pathways clustered into three functional categories: (*i*) immune and inflammatory responses, including cytokine-cytokine receptor interaction (hsa04060), chemokine-mediated signalling (GO:00700989), and Toll-like receptor signalling (hsa04620); (*ii*) pathogen-host interactions, encompassing viral protein interactions with cytokine receptors (hsa04061) and antiviral defence responses (GO:0051607); and (*iii*) cellular stress and pathological processes, including lipid metabolism in atherosclerosis (hsa05417) and transcriptional dysregulation in cancer (hsa05202). Across all shared pathways, B/Tunis864 maintained consistently higher enrichment scores than A/2497 (Table 1-2, Fig. S7a-d).

At 24 hpi, A/2497 engaged pathways involved in cellular maintenance and energy metabolism including oxidative phosphorylation (hsa00190), ribosomal function (hsa03010), and protein export machinery (hsa03060), suggesting a focus on basic cellular homeostasis (Table 1-2, Fig. S7e, Fig. S7g). In contrast, B/Tunis864 activated pathways related to immune response, spanning both innate and adaptive immunity including intestinal IgA production networks (hsa04672), antigen processing and presentation (hsa04612), NF-κB signalling (hsa04064), JAK-STAT signalling (hsa04630), antiviral innate immunity (GO:0140374), cytokine production regulation (GO:0001819), anti-bacterial defence responses (GO:0050829), and TNF-mediated signalling (GO:0033209). Additionally, B/Tunis864 triggered cellular stress and death pathways, including apoptosis (hsa04210) and ERK1/ERK2 cascade activation (GO:0070374) (Table 1-2, Fig. S7f, Fig. S7h).

### Pathway responses in THP-1 cells

At 4 hpi in THP-1 cells, pathway enrichment analysis revealed predominantly shared responses between strains, with 18 GO BP pathways commonly enriched by both strains, whilst A/2497 uniquely enriched 8 pathways and B/Tunis864 enriched 5 pathways (Table 2-1, Fig. 4e). Amongst KEGG pathways, 15 were shared between strains, with 5 unique to A/2497 and 3 to B/Tunis864 (Table 2-1, Fig. 4f). The shared core pathways clustered into two main functional categories: (*i*) immune and inflammatory responses, encompassing TNF signalling (hsa04668), inflammatory response (GO:0006954), chemokine-mediated signalling (GO:0070098), and NF-κB signalling (hsa04064); and (*ii*) pathogen-host interactions and cellular stress responses, including viral protein interactions with cytokine receptors (hsa04061), Notch signalling (GO:0007219), and homologous recombination-mediated DNA repair (GO:0000724) (Table 2-1, Fig. S8a-d). Notably, both strains triggered pathways involved in DNA regulation and repair processes.

**Table 2-1.** Pathways identified by Gene Set Enrichment Analysis (GO BP and KEGG) for HCjE cells inoculated with Ct strains A/2497 and B/Tunis864 at 4 hpi.

| ID | Description | enrichment score | enrichment score | enrichment score | enrichment score |
| --- | --- | --- | --- | --- | --- |
| **CtA\*** |  |  |  |  |  |
| GO:1902895 | positive regulation of miRNA transcription | 0.56 |  |  |  |
| GO:0007204 | positive regulation of cytosolic calcium ion concentration | 0.53 |  |  |  |
| GO:0098609 | cell-cell adhesion | 0.40 |  |  |  |
| GO:0001525 | angiogenesis | 0.37 |  |  |  |
| GO:0007267 | cell-cell signaling | 0.32 |  |  |  |
| GO:0051056 | regulation of small GTPase mediated signal transduction | -0.35 |  |  |  |
| GO:0006260 | DNA replication | -0.36 |  |  |  |
| GO:0000077 | DNA damage checkpoint signaling | -0.39 |  |  |  |
| hsa04512 | ECM-receptor interaction | 0.57 |  |  |  |
| hsa05213 | Endometrial cancer | 0.55 |  |  |  |
| hsa04024 | cAMP signaling pathway | 0.44 |  |  |  |
| hsa05163 | Human cytomegalovirus infection | 0.41 |  |  |  |
| hsa05200 | Pathways in cancer | 0.36 |  |  |  |
| **CtB\*** |  |  |  |  |  |
| GO:0042311 | vasodilation | 0.66 |  |  |  |
| GO:0034605 | cellular response to heat | 0.54 |  |  |  |
| GO:0019221 | cytokine-mediated signaling pathway | 0.46 |  |  |  |
| GO:0045944 | positive regulation of transcription by RNA polymerase II | 0.30 |  |  |  |
| GO:0031297 | replication fork processing | -0.63 |  |  |  |
| hsa04930 | Type II diabetes mellitus | 0.67 |  |  |  |
| hsa05120 | Epithelial cell signaling in Helicobacter pylori infection | 0.56 |  |  |  |
| hsa03430 | Mismatch repair | -0.71 |  |  |  |
| **Core\*** |  |  |  |  |  |
|  |  | **CtA** | **CtB** | **CtA_HIA** | **CtB_HIA** |
| GO:0002548 | monocyte chemotaxis | NA | 0.72 | 0.69 | 0.66 |
| GO:0070098 | chemokine-mediated signaling pathway | 0.74 | 0.71 | 0.76 | 0.62 |
| GO:0042531 | positive regulation of tyrosine phosphorylation of STAT protein | 0.67 | 0.71 | 0.69 | NA |
| GO:0003180 | aortic valve morphogenesis | 0.67 | 0.69 | 0.71 | NA |
| GO:0030593 | neutrophil chemotaxis | 0.70 | 0.65 | NA | NA |
| GO:0071347 | cellular response to interleukin-1 | 0.64 | 0.62 | 0.62 | 0.58 |
| GO:0071222 | cellular response to lipopolysaccharide | 0.52 | 0.56 | 0.52 | 0.50 |
| GO:0050729 | positive regulation of inflammatory response | NA | 0.54 | 0.56 | NA |
| GO:0071356 | cellular response to tumor necrosis factor | 0.53 | 0.52 | 0.56 | 0.49 |
| GO:0007219 | Notch signaling pathway | 0.44 | 0.42 | 0.41 | NA |
| GO:0006954 | inflammatory response | 0.41 | 0.42 | 0.42 | 0.42 |
| GO:0007186 | G protein-coupled receptor signaling pathway | 0.40 | 0.41 | 0.42 | 0.43 |
| GO:0008285 | negative regulation of cell population proliferation | 0.35 | 0.41 | 0.39 | 0.38 |
| GO:0043066 | negative regulation of apoptotic process | 0.35 | 0.36 | 0.35 | 0.37 |
| GO:0007155 | cell adhesion | 0.37 | 0.35 | 0.37 | 0.37 |
| GO:0010628 | positive regulation of gene expression | 0.33 | 0.33 | 0.35 | 0.32 |
| GO:0030335 | positive regulation of cell migration | 0.40 | NA | 0.43 | 0.41 |
| GO:0000724 | double-strand break repair via homologous recombination | -0.40 | -0.45 | NA | -0.46 |
| hsa04061 | Viral protein interaction with cytokine and cytokine receptor | 0.68 | 0.69 | 0.69 | 0.64 |
| hsa04060 | Cytokine-cytokine receptor interaction | 0.63 | 0.67 | 0.62 | 0.62 |
| hsa04668 | TNF signaling pathway | 0.59 | 0.61 | 0.63 | 0.62 |
| hsa05323 | Rheumatoid arthritis | 0.64 | 0.68 | 0.63 | 0.60 |
| hsa04062 | Chemokine signaling pathway | 0.50 | 0.46 | 0.46 | NA |
| hsa05144 | Malaria | 0.71 | 0.68 | 0.69 | NA |
| hsa05145 | Toxoplasmosis | 0.52 | 0.50 | 0.51 | NA |
| hsa04620 | Toll-like receptor signaling pathway | 0.51 | 0.54 | 0.51 | NA |
| hsa04625 | C-type lectin receptor signaling pathway | 0.52 | 0.54 | 0.55 | 0.54 |
| hsa05134 | Legionellosis | NA | 0.63 | 0.59 | NA |
| hsa05146 | Amoebiasis | 0.57 | 0.57 | 0.55 | 0.55 |
| hsa04657 | IL-17 signaling pathway | 0.55 | 0.58 | 0.54 | 0.55 |
| hsa05417 | Lipid and atherosclerosis | 0.40 | 0.46 | 0.42 | NA |
| hsa04064 | NF-kappa B signaling pathway | 0.57 | 0.57 | 0.54 | 0.53 |
| hsa04630 | JAK-STAT signaling pathway | 0.53 | 0.55 | 0.50 | 0.56 |
| **HIA<sup>†</sup>** |  |  |  |  |  |
|  |  |  |  | **CtA_HIA** | **CtB_HIA** |
| GO:0001974 | blood vessel remodeling |  |  | 0.68 | NA |
| GO:0045600 | positive regulation of fat cell differentiation |  |  | 0.63 | NA |
| GO:0043616 | keratinocyte proliferation |  |  | 0.60 | NA |
| GO:0030216 | keratinocyte differentiation |  |  | 0.58 | NA |
| GO:0001938 | positive regulation of endothelial cell proliferation |  |  | 0.55 | NA |
| GO:0014032 | neural crest cell development |  |  | 0.55 | NA |
| GO:0048661 | positive regulation of smooth muscle cell proliferation |  |  | 0.52 | NA |
| GO:0008284 | positive regulation of cell population proliferation |  |  | 0.37 | NA |

**Table 2-2.**
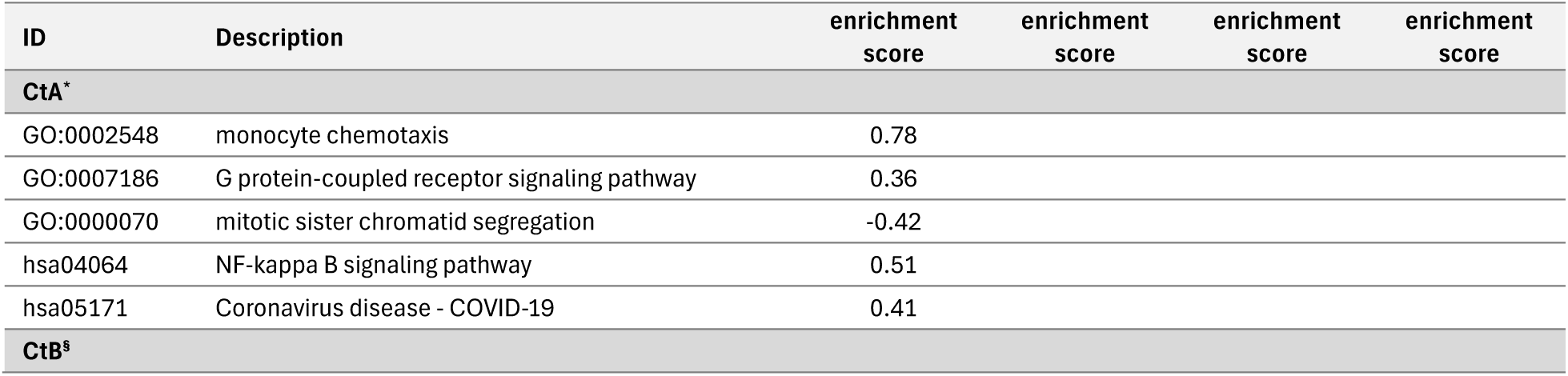
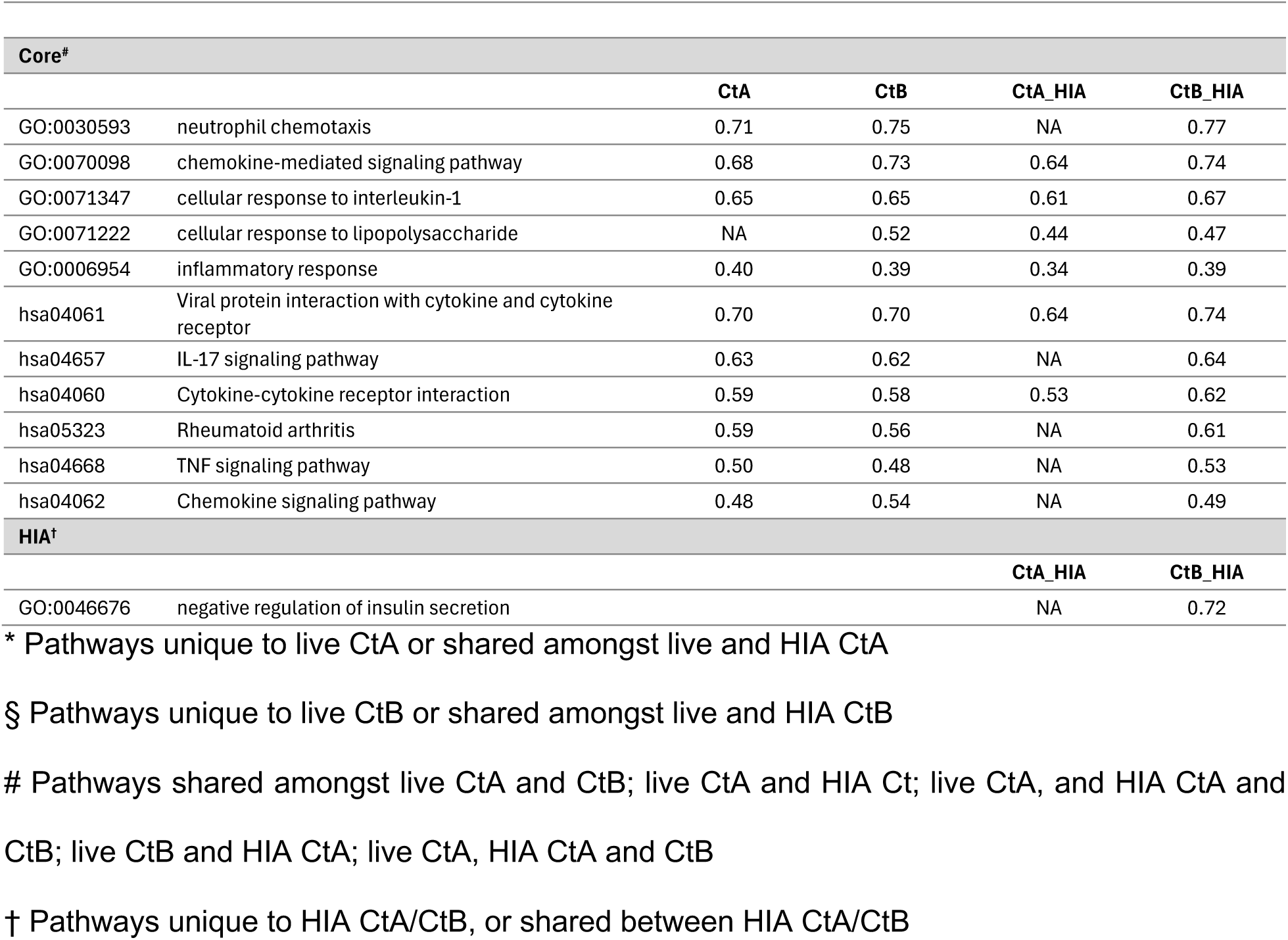
Pathways identified by Gene Set Enrichment Analysis (GO BP and KEGG) for HCjE cells inoculated with Ct strains A/2497 and B/Tunis864 at 24 hpi.

| ID | Description | enrichment score | enrichment score | enrichment score | enrichment score |
| --- | --- | --- | --- | --- | --- |
| **CtA<sup>\*</sup>** |  |  |  |  |  |
| GO:0002548 | monocyte chemotaxis | 0.78 |  |  |  |
| GO:0007186 | G protein-coupled receptor signaling pathway | 0.36 |  |  |  |
| GO:0000070 | mitotic sister chromatid segregation | -0.42 |  |  |  |
| hsa04064 | NF-kappa B signaling pathway | 0.51 |  |  |  |
| hsa05171 | Coronavirus disease - COVID-19 | 0.41 |  |  |  |
| **CtB<sup>§</sup>** |  |  |  |  |  |

| Core<sup>#</sup> |  | CtA | CtB | CtA_HIA | CtB_HIA |
| --- | --- | --- | --- | --- | --- |
| GO:0030593 | neutrophil chemotaxis | 0.71 | 0.75 | NA | 0.77 |
| GO:0070098 | chemokine-mediated signaling pathway | 0.68 | 0.73 | 0.64 | 0.74 |
| GO:0071347 | cellular response to interleukin-1 | 0.65 | 0.65 | 0.61 | 0.67 |
| GO:0071222 | cellular response to lipopolysaccharide | NA | 0.52 | 0.44 | 0.47 |
| GO:0006954 | inflammatory response | 0.40 | 0.39 | 0.34 | 0.39 |
| hsa04061 | Viral protein interaction with cytokine and cytokine receptor | 0.70 | 0.70 | 0.64 | 0.74 |
| hsa04657 | IL-17 signaling pathway | 0.63 | 0.62 | NA | 0.64 |
| hsa04060 | Cytokine-cytokine receptor interaction | 0.59 | 0.58 | 0.53 | 0.62 |
| hsa05323 | Rheumatoid arthritis | 0.59 | 0.56 | NA | 0.61 |
| hsa04668 | TNF signaling pathway | 0.50 | 0.48 | NA | 0.53 |
| hsa04062 | Chemokine signaling pathway | 0.48 | 0.54 | NA | 0.49 |
| HIA<sup>†</sup> |  |  |  | CtA_HIA | CtB_HIA |
| GO:0046676 | negative regulation of insulin secretion |  |  | NA | 0.72 |

\* Pathways unique to live CtA or shared amongst live and HIA CtA
§ Pathways unique to live CtB or shared amongst live and HIA CtB

At 4 hpi, A/2497 specifically triggered DNA replication machinery (GO:0006260) and DNA damage checkpoint signalling (GO:0000077), alongside cellular communication pathways including cell-cell signalling (GO:0007267) and cAMP signalling (hsa04024) (Table 2-1, Fig. S8e, Fig. S8g). In contrast, B/Tunis864 triggered DNA mismatch repair (hsa03430) and replication fork processing (GO:0031297) pathways (Table 2-1, Fig. S8f, Fig. S8h).

By 24 hpi, pathway enrichment had diminished, with only 5 GO BP pathways commonly enriched by both strains and 3 unique to A/2497, whilst no GO BP pathways were identified specifically for B/Tunis864 (Table 2-2, Fig. 4g). Amongst KEGG pathways, 6 remained shared between strains, 2 were unique to A/2497, and no pathways were specific to B/Tunis864 (Table 2-2, Fig. 4h). The shared pathways at 24 hpi were predominantly inflammatory and tissue-damaging responses, including TNF signalling (hsa04668), neutrophil chemotaxis (GO:0030593), inflammatory response (GO:0006954), chemokine-mediated signalling (GO:0070098), and rheumatoid arthritis-associated pathways (hsa05323) (Table 2-2, Fig. S9a-d). At 24 hpi, A/2497 maintained unique activation of immune-regulatory pathways, specifically monocyte chemotaxis (GO:0002548) and NF-κB signalling (hsa04064), alongside a cellular process pathway involving mitotic sister chromatid segregation (GO:0000070) (Table 2-2, Fig. S9e, Fig. S9f).

### Pathway-associated genes in KEGG and GO BP enrichment

In HCjE cells at 4 hpi, pathway enrichment analysis identified 89 and 139 genes associated with significantly enriched pathways for A/2497 and B/Tunis864 infection, respectively. Of the 181 unique pathway-associated genes examined, 35 exhibited significant differential expression relative to uninfected controls (adjusted *P* < 0.05) (Fig. 5a, Fig. S6i). Expression pattern analysis revealed predominantly balanced responses between strains: 16 genes (45.7%) demonstrated concordant upregulation, whilst 7 were CtA-specific and 8 were CtB-specific. Additionally, 3 genes showed CtB-dominant responses, and 1 gene displayed opposite regulatory directions (Fig. 5b). Notable CtA-specific inflammatory mediators included IL1RN, CSF1R, IL20RB, and S100A9. In contrast, CtB-specific/dominant genes encompassed critical NF-κB pathway regulators (NFKBIZ, NFKBIA, ZC3H12A) and pro-inflammatory mediators (IL6, CXCL1, CXCL8, NR3C1) (Fig. 5b).

**Fig. 5.**
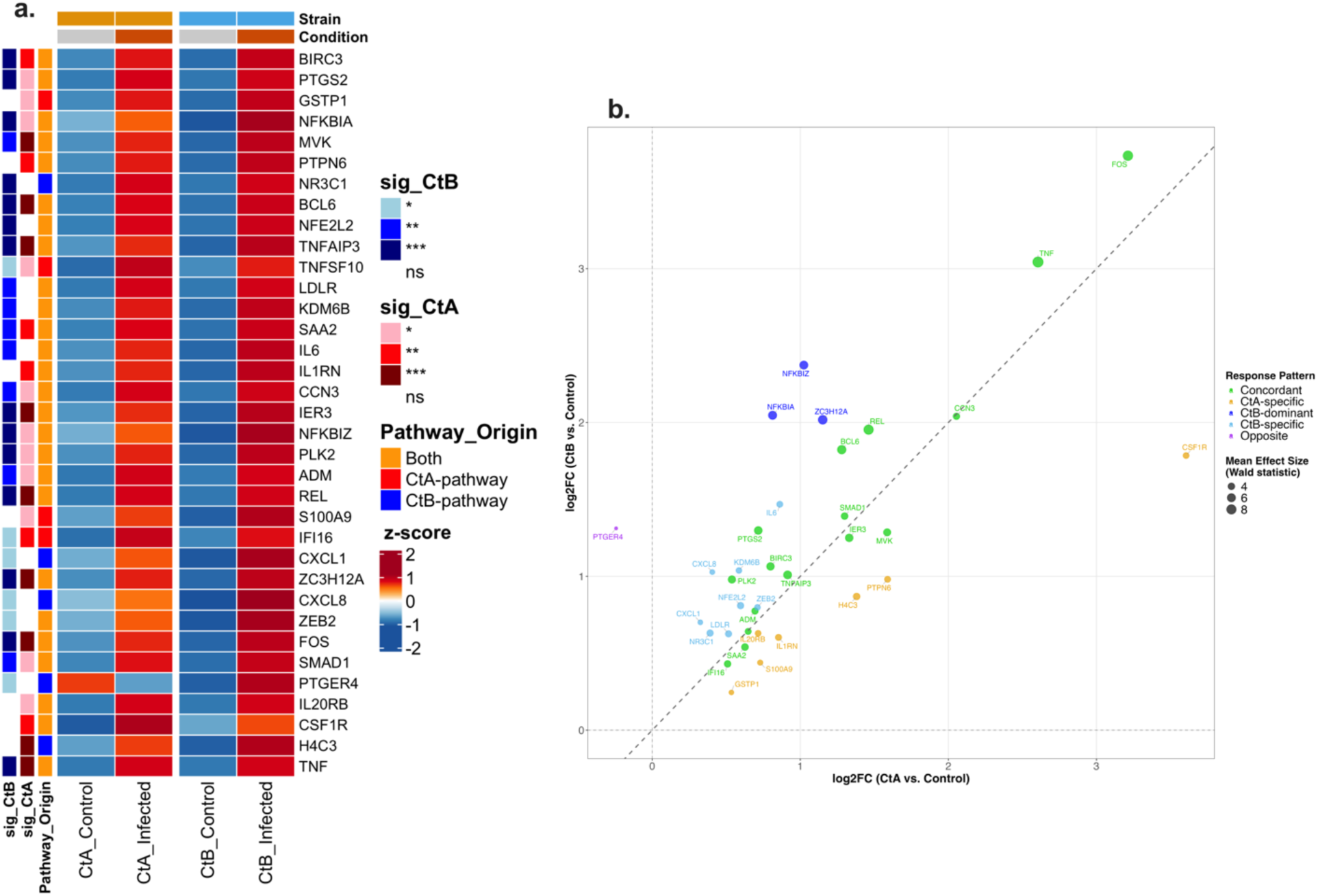
Differential expression patterns of pathway-associated genes in HCjE cells infected with Ct strains at 4 hpi. (**a**) Heatmap showing normalised expression (Z-scores) of pathway-associated genes significantly differentially expressed (adjusted *P* < 0.05) in CtA (A/2497) and/or CtB (B/Tunis864) infection. Columns: mean expression across replicates for controls and infected samples. Rows: genes ordered by pathway origin. Row annotations: pathway source (red: CtA-pathway; blue: CtB-pathway; orange: both; grey: neither) and strain-specific significance levels (*** *P* < 0.001, ** *P* < 0.01, * *P* < 0.05, white: ns). Column annotations: condition (grey: control; orange: infected) and strain (orange: CtA; blue: CtB). (**b**) Scatter plot of log2FC values comparing CtA *vs*. CtB responses for significantly differentially expressed genes. Points coloured by response pattern: orange (CtA-specific), light blue (CtB-specific), green (Concordant, |Δlog2FC| < 0.585), red (CtA-dominant, ≥ 1.5-fold stronger), dark blue (CtB-dominant, ≥ 1.5-fold stronger), purple (Opposite directions). Point size reflects mean effect size (Wald statistic). Diagonal dashed line indicates equal responses; axes intersect at baseline. Top genes per response pattern are labelled.

At 24 hpi in HCjE cells, pathway enrichment analysis identified 29 and 163 genes associated with significantly enriched pathways for A/2497 and B/Tunis864 infection, respectively. Of the 172 unique pathway-associated genes examined, 86 exhibited significant differential expression (Fig. 6a, Fig. S7i). Expression patterns revealed markedly strain-differentiated responses: B/Tunis864 infection induced 54 strain-specific and 11 strain-dominant genes (collectively 75.6% of significant genes), whilst A/2497 elicited only 1 strain-specific gene. Concordant responses declined to 4 genes, and 16 genes displayed opposite regulatory directions (Fig. 6b). The B/Tunis864-dominant response was characterised by robust interferon-stimulated gene (ISG) activation, including IFIT1, IFIT2, IFIT3, HERC5, MX1, MX2, OAS1, OAS2, and RSAD2, alongside interferon-inducible chemokines (CXCL10, CXCL11) and pro-inflammatory mediators (IL1A, IL6, CCL3, CCL5, TNFSF10, TNFSF13B). STAT1, a key interferon signalling transcription factor, was also exclusively upregulated in B/Tunis864 infection. ISG15 was significantly upregulated in CtB infection while there were no significant changes in CtA infection further highlighting divergent immune responses between strains (Fig. 6b).

**Fig. 6.**
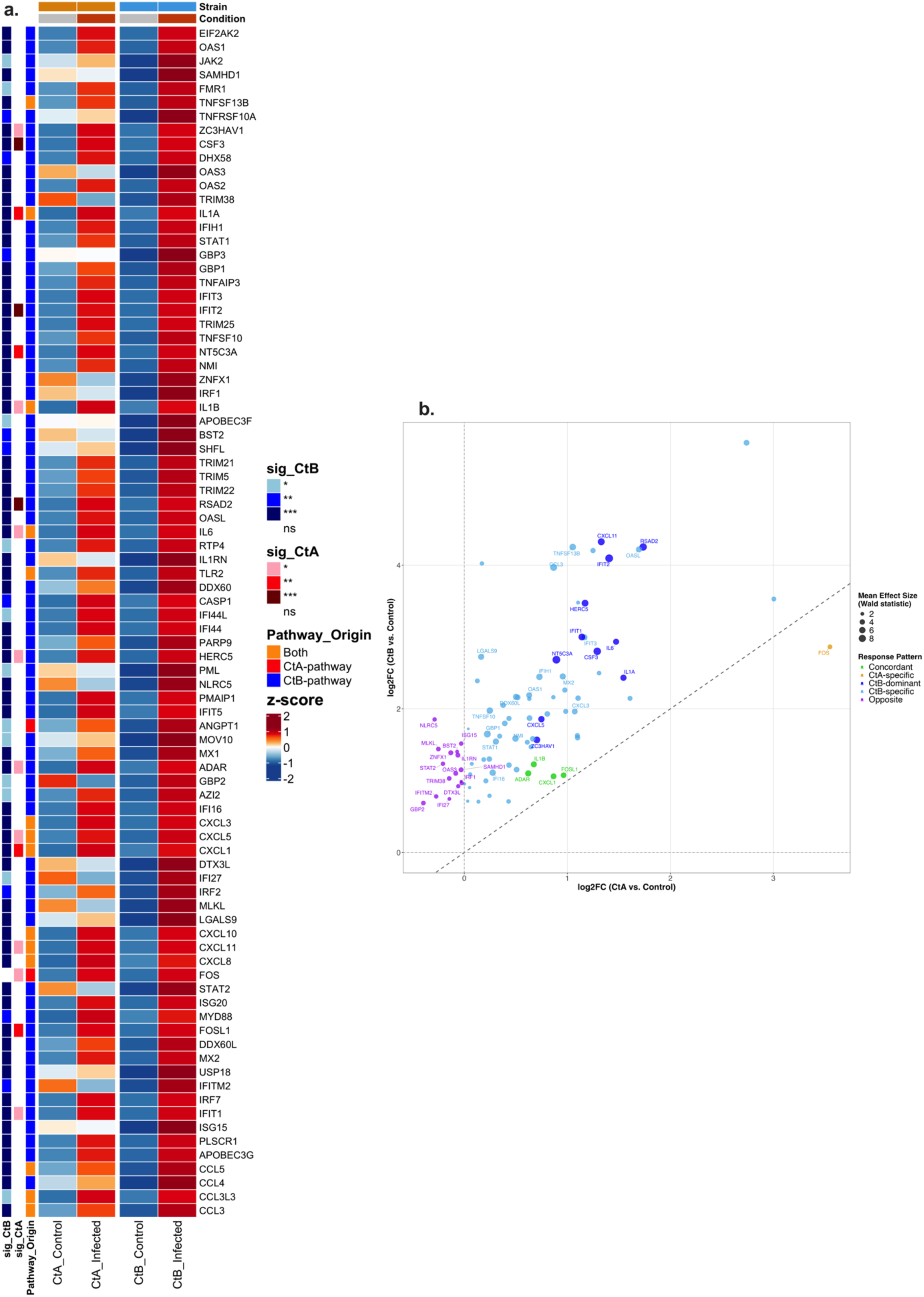
Differential expression patterns of pathway-associated genes in HCjE cells infected with Ct strains at 24 hpi. (**a**) Heatmap showing normalised expression (Z-scores) of pathway-associated genes significantly differentially expressed (adjusted *P* < 0.05) in CtA (A/2497) and/or CtB (B/Tunis864) infection. Columns: mean expression across replicates for controls and infected samples. Rows: genes ordered by pathway origin. Row annotations: pathway source (red: CtA-pathway; blue: CtB-pathway; orange: both; grey: neither) and strain-specific significance levels (*** *P* < 0.001, ** *P* < 0.01, * *P* < 0.05, white: ns). Column annotations: condition (grey: control; orange: infected) and strain (orange: CtA; blue: CtB). (**b**) Scatter plot of log2FC values comparing CtA *vs*. CtB responses for significantly differentially expressed genes. Points coloured by response pattern: orange (CtA-specific), light blue (CtB-specific), green (Concordant, |Δlog2FC| < 0.585), red (CtA-dominant, ≥ 1.5-fold stronger), dark blue (CtB-dominant, ≥ 1.5-fold stronger), purple (Opposite directions). Point size reflects mean effect size (Wald statistic). Diagonal dashed line indicates equal responses; axes intersect at baseline. Top genes per response pattern are labelled.

In THP-1 cells at 4 hpi, pathway enrichment analysis identified 225 and 136 genes associated with significantly enriched pathways for A/2497 and B/Tunis864 infection, respectively. Of the 305 unique pathway-associated genes examined, 132 exhibited significant differential expression (Fig. 7a, Fig. S8i). Expression analysis revealed predominantly concordant inflammatory responses characteristic: 76 genes (57.6%) demonstrated similar upregulation patterns between strains. These included pro-inflammatory cytokines (TNF, IL1B, CSF3), NF-κB pathway components (NFKBIZ, REL, NFKBIA, NFKB1), and chemokines (CXCL1, CXCL2, CXCL3, CXCL8, CCL3, CCL3L3, CCL5). Modest strain-biased responses were observed: 36 genes were CtA-specific and 8 were CtA-dominant (including CCL4, CCL20, CSF2, TNIP3), whilst 10 genes were CtB-specific and 1 was CtB-dominant. Only 1 gene displayed opposite regulation (Fig. 7b).

**Fig. 7.**
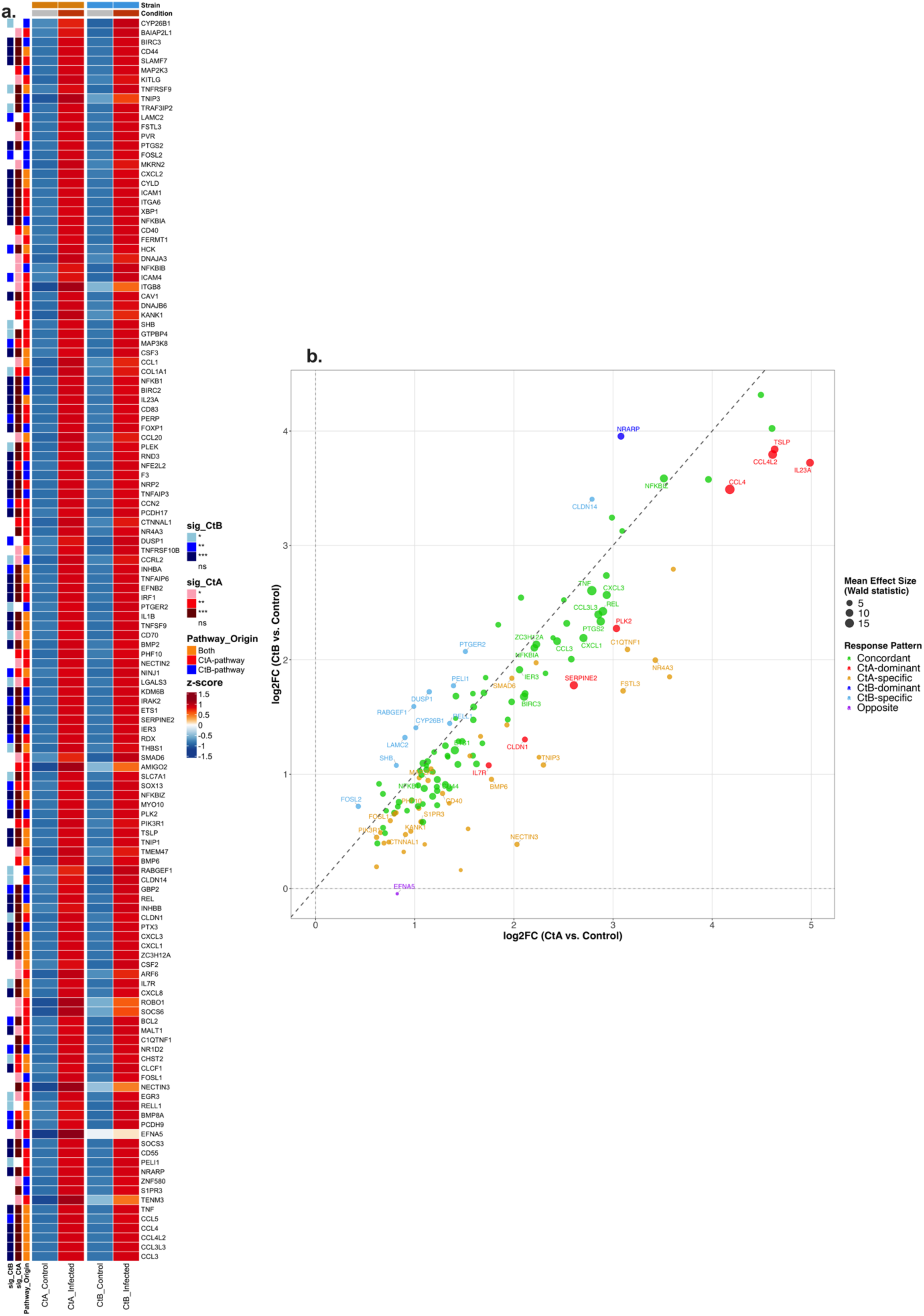
Differential expression patterns of pathway-associated genes in THP-1 cells infected with Ct strains at 4 hpi. (**a**) Heatmap showing normalised expression (Z-scores) of pathway-associated genes significantly differentially expressed (adjusted *P* < 0.05) in CtA (A/2497) and/or CtB (B/Tunis864) infection. Columns: mean expression across replicates for controls and infected samples. Rows: genes ordered by pathway origin. Row annotations: pathway source (red: CtA-pathway; blue: CtB-pathway; orange: both; grey: neither) and strain-specific significance levels (*** *P* < 0.001, ** *P* < 0.01, * *P* < 0.05, white: ns). Column annotations: condition (grey: control; orange: infected) and strain (orange: CtA; blue: CtB). (**b**) Scatter plot of log2FC values comparing CtA *vs*. CtB responses for significantly differentially expressed genes. Points coloured by response pattern: orange (CtA-specific), light blue (CtB-specific), green (Concordant, |Δlog2FC| < 0.585), red (CtA-dominant, ≥ 1.5-fold stronger), dark blue (CtB-dominant, ≥ 1.5-fold stronger), purple (Opposite directions). Point size reflects mean effect size (Wald statistic). Diagonal dashed line indicates equal responses; axes intersect at baseline. Top genes per response pattern are labelled.

At 24 hpi in THP-1 cells, pathway enrichment analysis identified 51 and 45 genes associated with significantly enriched pathways for A/2497 and B/Tunis864 infection, respectively. Of the 58 unique pathway-associated genes examined, 28 exhibited significant differential expression (Fig. 8a, Fig. S9g). Expression patterns revealed sustained concordant responses: 19 genes (67.9%) maintained similar upregulation between strains, demonstrating persistent inflammatory activation. Concordant genes included chemokines (CCL1, CCL3L3, CCL4, CCL4L2, CCL5, CXCL3, CXCL5, CXCL8), pro-inflammatory cytokines (IL1B, IL23A, CSF3), and TNF superfamily members (TNFSF15, TNFAIP6). Minimal strain-differentiation was observed: 5 genes were CtA-specific (including BMP6, CD70, TNFRSF10A) and 3 were CtA-dominant (TNF, CXCL1, CCL3), whilst only 1 gene was CtB-specific (IL32) (Fig. 8b).

**Fig. 8.**
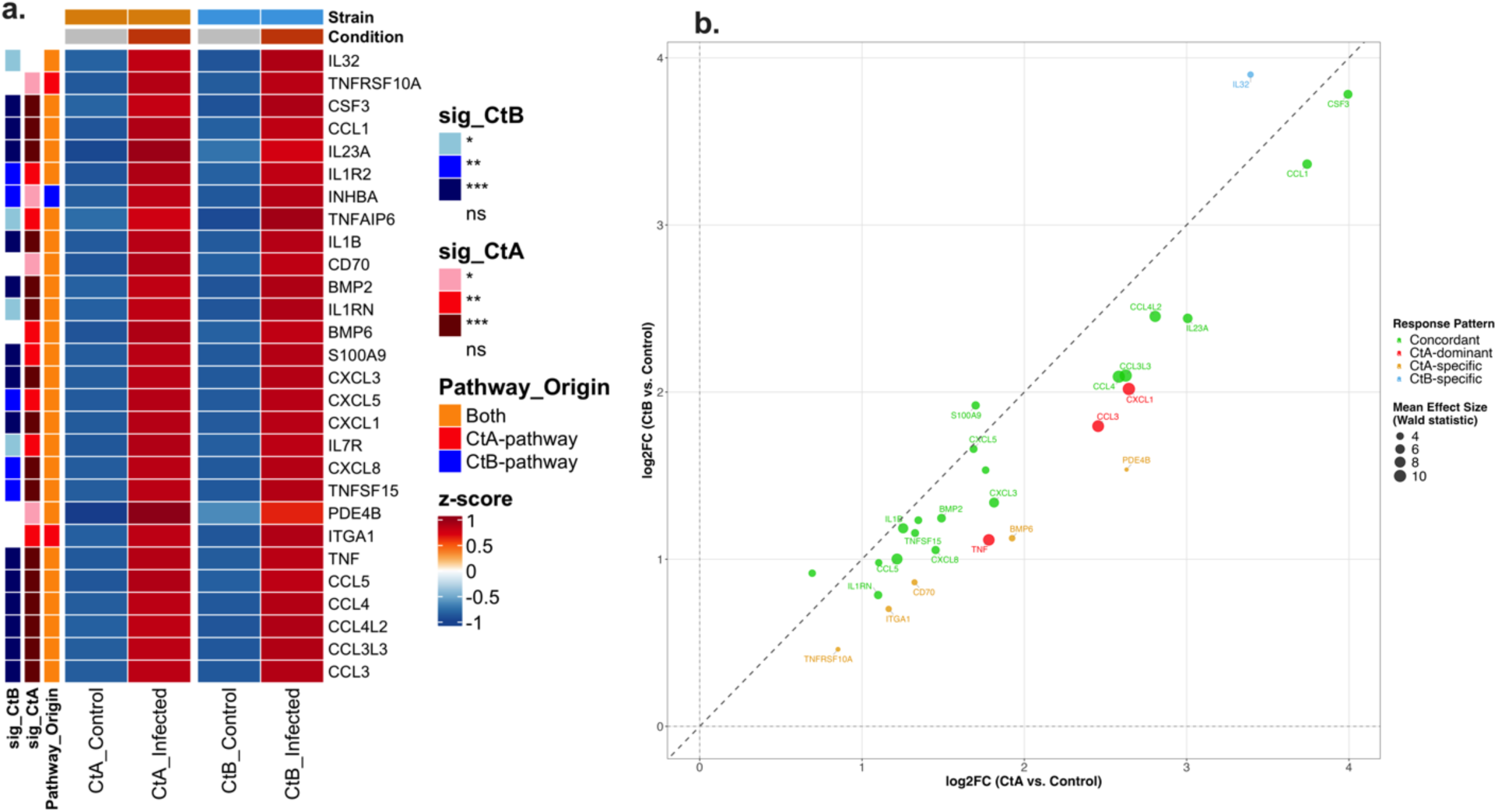
Differential expression patterns of pathway-associated genes in THP-1 cells infected with Ct strains at 24 hpi. (**a**) Heatmap showing normalised expression (Z-scores) of pathway-associated genes significantly differentially expressed (adjusted *P* < 0.05) in CtA (A/2497) and/or CtB (B/Tunis864) infection. Columns: mean expression across replicates for controls and infected samples. Rows: genes ordered by pathway origin. Row annotations: pathway source (red: CtA-pathway; blue: CtB-pathway; orange: both; grey: neither) and strain-specific significance levels (*** *P* < 0.001, ** *P* < 0.01, * *P* < 0.05, white: ns). Column annotations: condition (grey: control; orange: infected) and strain (orange: CtA; blue: CtB). (**b**) Scatter plot of log2FC values comparing CtA *vs*. CtB responses for significantly differentially expressed genes. Points coloured by response pattern: orange (CtA-specific), light blue (CtB-specific), green (Concordant, |Δlog2FC| < 0.585), red (CtA-dominant, ≥ 1.5-fold stronger), dark blue (CtB-dominant, ≥ 1.5-fold stronger), purple (Opposite directions). Point size reflects mean effect size (Wald statistic). Diagonal dashed line indicates equal responses; axes intersect at baseline. Top genes per response pattern are labelled.

## Discussion

In this study, we characterised the shared and distinct transcriptional responses of Ct strains A/2497 and B/Tunis864 across early- and mid-developmental stages (4 and 24 hpi) induced in HCjE and THP-1 cells. The kinetics of responses to Ct infection differed between cell types: THP-1 cells showed peak transcriptional activation at 4 hpi, whereas HCjE cells demonstrated peak transcriptional activation at 24 hpi. HCjE cells exhibited strain related transcriptional responses, contrasting with THP-1 cells that exhibited a response characterised by overlapping DEGs and enriched pathways regardless of Ct strain.

In HCjE cells, the number of DEGs and enriched pathways progressively increased from 4 to 24 hpi, consistent with established patterns of epithelial cell adaptation during chlamydial infection [27,52]. This temporal transcriptomic activation mirrors recent chromatin accessibility studies in Hep-2 cells, which demonstrated dramatic increases from 864 differentially accessible regions at early time points to over 3,000 regions by 48 hpi, reflecting Ct’s progressive exploitation of epithelial cellular machinery throughout its developmental cycle [8,23,52]. Recent studies have shown that HCjE cells support replication of major Ct genovars whilst exhibiting genovar-specific cytokine production patterns [53–55], consistent with our observation of marked strain-specific transcriptional differences where CtB consistently elicited stronger responses than CtA at both time points.

Conversely, THP-1 macrophages exhibited decreasing pathway enrichment over time, consistent with the temporal dynamics of macrophage responses to intracellular pathogens [56]. This pattern likely reflects the binary outcome of macrophage-Chlamydia interactions: either successful pathogen clearance through robust early inflammatory responses, or establishment of infection through evasion of macrophage defence mechanisms, including subversion of phagosome maturation, modulation of pro-inflammatory cytokine production, and interference with macrophage polarisation [57–63]. Unlike the pronounced strain-specific responses observed in HCjE cells, THP-1 macrophages displayed predominantly shared responses to CtA and CtB, suggesting that macrophages deploy a core response regardless of infecting strain. This pattern of strain-invariant macrophage responses has been similarly observed in infections with *Mycobacterium abscessus* Smooth and Rough variants [64], suggesting that macrophages may prioritise rapid, generalised defence mechanisms over strain-specific responses during initial encounter with Ct.

At 4 hpi, HCjE cells displayed markedly different transcriptional responses to CtA and CtB infection, with CtB inducing nearly 3-fold more DEGs than CtA. Despite activating distinct strain-specific pathways, both strains upregulated core stress responses, including integrated stress response signalling (GO:0140467) and inflammatory response (GO:0006954), with CtB consistently exhibiting higher enrichment scores across these shared pathways. By 24 hpi, both the number of DEGs and enriched pathways increased substantially compared to 4 hpi, consistent with prior studies in epithelial cells demonstrating that immune response activation to Ct becomes pronounced during mid-cycle infection (6-24 hpi) [24,65].

At 24 hpi, CtA activated pathways associated with cellular maintenance and metabolic homeostasis in HCjE cells, including oxidative phosphorylation (hsa00190), ribosomal function (hsa03010), and protein export machinery (hsa03060). This metabolic signature represents hijacking of host cellular resources to support bacterial ATP requirements through both glucose-6-phosphate utilisation and direct ATP scavenging via nucleotide transporters [58,66,67]. In contrast, CtB mounted a robust immune and stress response. The extensive transcriptional activation induced by CtB in HCjE cells, particularly the pronounced ISG response at 24 hpi (CXCL10, CXCL11, MX1, OAS2, ISG20, TRIM5, TRIM22, STAT1), may explain the prior observations on more severe pathology associated with CtB than CtA in trachoma patients [36]. This ISG signature reflects type-I IFN pathway activation through STING-mediated recognition of chlamydial cyclic di-AMP, a response that, whilst initially protective, may contribute to the immunopathology characteristic of trachoma when dysregulated [53,68–72]. CtB induced a pronounced type II interferon response compared to CtA, suggesting that CtA may have evolved specific mechanisms to evade IFN-γ-mediated tryptophan starvation through indoleamine 2,3-dioxygenase (IDO) inhibition [73–76], consistent with broader *Chlamydia* immune evasion strategies that include suppression of cytokine and chemokine production by epithelial cells [77,78] and interference with antigen presentation through downregulation of MHC class I and II molecules on antigen-presenting cells [77,79–81]. Furthermore, CtB’s selective activation of cell death pathways, including apoptosis (hsa04210) and pyroptosis (GO:0070269), alongside adaptive immune response pathways such as antigen processing and presentation (hsa04612) and JAK-STAT signalling (hsa04630), indicate an intense inflammatory infection course that likely contributes to the tissue damage in trachoma [82–85].

In THP-1 macrophages, CtA and CtB infection induced substantial overlap in both genes and pathways, clustering predominantly into immune and inflammatory responses, pathogen-host interactions, and tissue remodelling processes. However, despite this shared core response, CtA consistently elicited more pronounced transcriptional activation than CtB at both time points. At 4 hpi, both strains induced early activation of DNA damage response pathways, with CtA specifically upregulating DNA replication machinery and damage checkpoint signalling, whilst CtB preferentially activated DNA mismatch repair pathways. These distinct pathway signatures suggest differential strategies for managing Ct-induced genomic stress [23]. Recent findings have demonstrated that Ct disrupts homologous recombination repair through PP2A-mediated inactivation of ataxia telangiectasia mutated (ATM) kinase [86], providing a molecular basis for the observed DNA damage responses.

By 24 hpi, CtA stimulated higher overall transcriptomic activity than CtB, uniquely activating key immune regulatory pathways including monocyte chemotaxis (GO:0002548) and NF-κB signalling (hsa04064), indicating sustained inflammatory programming. CtA’s specific activation of cellular chemotaxis and cAMP signalling pathways suggests enhanced capacity to modulate macrophage function. Three dominant CtA-induced genes identified in our analysis including CCL5, DNAJB6, and TNIP3 which play crucial roles in immune regulation: CCL5 mediates protective Th1 immunity through the CCR5-CCL5 axis, whilst TNIP3 functions as a negative regulator of NF-κB signalling [87,88]. Notably, enhanced CCL5 expression has been observed in conjunctival swabs from Gambian children with active trachoma compared to healthy controls [89]. The differential activation of NF-κB signalling pathways between CtA and CtB aligns with previous studies demonstrating functionally distinct NF-κB regulatory strategies amongst Chlamydia species [90,91], and may represent a key mechanism underlying the varied clinical outcomes associated with different Ct genovars [77,87,92]. The progressive reduction in THP-1 responsiveness observed from 4 to 24 hpi likely reflects temporal modulation corresponding to the chlamydial developmental cycle: robust immune activation during EB to RB conversion, followed by immune resolution during RB replication phases [93,94]. This temporal attenuation is mediated by sophisticated bacterial immune evasion strategies [95], likely through CPAF, which progressively cleaves the p65/RelA transcription factor, thereby reducing NF-κB signalling and attenuating IL-1β-dependent IL-8 secretion [94,96,97].

This study has several limitations. First, whilst PCA effectively captured sources of variation, the relatively modest variance explained by PC1 and PC2 suggests that additional biological or technical factors may contribute to the observed gene expression patterns. Second, we examined only two discrete time points (4 and 24 hpi), which, whilst capturing early- and mid-cycle responses, precludes a comprehensive understanding of the complete temporal dynamics of host-Ct interactions. Incorporation of additional time points, particularly during late-cycle RB to EB differentiation (>40 hpi), would provide a more complete picture of the infection trajectory. Third, the functional interpretation relied exclusively on computational enrichment analysis of GO and KEGG pathways, which, whilst informative, may not capture all relevant biological processes or reflect actual protein-level changes. Finally, whilst this study identified distinct pathway activation patterns between CtA and CtB, it did not investigate the underlying molecular mechanisms that drive these genovar-specific host responses.

Given that the conjunctival epithelium represents the primary target of ocular Ct infection, the abundance of genovar-specific DEGs in epithelial cells may suggest that these cells serve as the principal site where distinct host responses to different Ct genovars are established. Our findings suggest that cell type determines fundamental response architecture whilst genovar specificity may fine-tune pathogenic outcomes. The progressive pathway enrichment in epithelial cells reflects mounting host immune and inflammatory responses to infection, alongside bacterial hijacking of cellular metabolic machinery to support replication, whereas the peak-then-decline response pattern observed in macrophages reflects the transition from an early strong response to the infection to a progressively subverted response during intracellular replication of Ct. The pronounced interferon-stimulated gene signature associated with CtB infection, particularly the upregulation of CXCL10, CXCL11, and STAT1, may offer potential biomarkers for monitoring disease severity and inflammatory responses that correlate with scarring progression in trachoma. The robust inflammatory signatures induced by CtB in HCjE cells provide molecular support for clinical observations linking CtB with more severe trachomatous disease, thereby bridging these findings with epidemiological findings. Collectively, these findings support genovar-informed interventions, such as prioritising mass drug administration based on circulating Ct genovars in trachoma-endemic populations, potentially optimising resource allocation and accelerating progress towards the WHO 2030 elimination goals.

## Acknowledgments

We would like to thank the Bioimaging Facility of the London School of Hygiene and Tropical Medicine for their technical assistance throughout the imaging steps. We extend our thanks to Dr Robert Butcher and Dr Ronan Doyle for their insightful guidance on the design of the study and data analysis.

## Author contributions

Conceived study: EG, MJH. Conducted laboratory analyses: EG, MJH. Conducted data analysis and interpretation: EG. Drafted manuscript: EG, MJH. Commented, edited, approved manuscript: EG, MJH.

## Declarations

### Ethics approval and consent to participate

Not applicable.

### Availability of data and materials

Sequencing data in the form of fastq.gz files used in this study can be accessed from the National Center for Biotechnology Information (NCBI) project accession PRJNA1133641 (Accession SAMN42382715-SAMN42382774).

