## Supplementary Data for "Comparative transcriptomic profiling of human conjunctival epithelial cells and macrophages in response to *Chlamydia trachomatis* genovars A and B in early- and mid-infection cycles"

**Table S1.** Read trimming statistics per sample using Cutadapt.

| Sample | Number of input reads [Millions] | Average input read length | Number of output reads [Millions] | Average output read length |
|---|---|---|---|---|
| HCjE_CTA_4_1 | 5.62 | 101 | 5.44 | 78.43 |
| HCjE_CTA_4_2 | 3.85 | 101 | 3.67 | 73.55 |
| HCjE_CTA_4_3 | 3.85 | 101 | 3.68 | 75.1 |
| HCjE_CTA_24_1 | 6.96 | 101 | 6.7 | 79.67 |
| HCjE_CTA_24_2 | 4.64 | 101 | 4.47 | 80.26 |
| HCjE_CTA_24_3 | 4.17 | 101 | 4 | 81.75 |
| HCjE_CTA_HIA_4_1 | 5.16 | 101 | 5.01 | 82.48 |
| HCjE_CTA_HIA_4_2 | 3.35 | 101 | 3.25 | 82.92 |
| HCjE_CTA_HIA_4_3 | 1.28 | 101 | 1.16 | 68.43 |
| HCjE_CTA_HIA_24_1 | 5.27 | 101 | 5.05 | 78.84 |
| HCjE_CTA_HIA_24_2 | 4.16 | 101 | 4 | 79.1 |
| HCjE_CTA_HIA_24_3 | 5.3 | 101 | 4.99 | 77.32 |
| HCjE_CTB_4_1 | 5.14 | 101 | 4.98 | 81.68 |
| HCjE_CTB_4_2 | 4.89 | 101 | 4.73 | 80.01 |
| HCjE_CTB_4_3 | 3.27 | 101 | 3.13 | 78.46 |
| HCjE_CTB_24_1 | 4.72 | 101 | 4.56 | 78.49 |
| HCjE_CTB_24_2 | 4.86 | 101 | 4.7 | 82.64 |
| HCjE_CTB_24_3 | 4.58 | 101 | 4.4 | 76.93 |
| HCjE_CTB_HIA_4_1 | 6.11 | 101 | 5.87 | 79.79 |
| HCjE_CTB_HIA_4_2 | 3.6 | 101 | 3.48 | 79.38 |
| HCjE_CTB_HIA_4_3 | 3.96 | 101 | 3.84 | 81.08 |
| HCjE_CTB_HIA_24_1 | 4.26 | 101 | 4.13 | 78.79 |
| HCjE_CTB_HIA_24_2 | 4.2 | 101 | 4.05 | 77.19 |
| HCjE_CTB_HIA_24_3 | 6.4 | 101 | 6.19 | 80.3 |
| HCjE_Con_4_1 | 7.12 | 101 | 6.86 | 81.9 |
| HCjE_Con_4_2 | 6.45 | 101 | 6.22 | 83.17 |
| HCjE_Con_4_3 | 3.64 | 101 | 3.5 | 81.29 |
| HCjE_Con_24_1 | 4.41 | 101 | 4.27 | 77.75 |
| HCjE_Con_24_2 | 3.42 | 101 | 3.27 | 76.52 |
| HCjE_Con_24_3 | 3.47 | 101 | 3.26 | 73.13 |
| THP1_CTA_4_1 | 4.49 | 101 | 4.32 | 79.3 |
| THP1_CTA_4_2 | 3.89 | 101 | 3.77 | 81.85 |
| THP1_CTA_4_3 | 4 | 101 | 3.9 | 83.23 |
| THP1_CTA_24_1 | 5.6 | 101 | 5.39 | 81.38 |
| THP1_CTA_24_2 | 4 | 101 | 3.84 | 80.88 |
| THP1_CTA_24_3 | 3.48 | 101 | 3.26 | 74.75 |
| THP1_CTA_HIA_4_1 | 3.45 | 101 | 3.35 | 83.34 |
| THP1_CTA_HIA_4_2 | 2.96 | 101 | 2.87 | 79.41 |
| THP1_CTA_HIA_4_3 | 4.78 | 101 | 4.55 | 76.86 |
| THP1_CTA_HIA_24_1 | 3.14 | 101 | 2.98 | 75.64 |

| | | | | |
|---|---|---|---|---|
| THP1_CTA_HIA_24_2 | 3.67 | 101 | 3.52 | 78.42 |
| THP1_CTA_HIA_24_3 | 6.17 | 101 | 5.98 | 81.02 |
| THP1_CTB_4_1 | 4.02 | 101 | 3.88 | 83.79 |
| THP1_CTB_4_2 | 4.61 | 101 | 4.43 | 78.79 |
| THP1_CTB_4_3 | 4.37 | 101 | 4.21 | 81.78 |
| THP1_CTB_24_1 | 3.47 | 101 | 3.36 | 81.65 |
| THP1_CTB_24_2 | 3.32 | 101 | 3.2 | 81.57 |
| THP1_CTB_24_3 | 4.59 | 101 | 4.4 | 79.35 |
| THP1_CTB_HIA_4_1 | 4.89 | 101 | 4.7 | 81.66 |
| THP1_CTB_HIA_4_2 | 3.99 | 101 | 3.77 | 74.57 |
| THP1_CTB_HIA_4_3 | 4.08 | 101 | 3.93 | 80.75 |
| THP1_CTB_HIA_24_1 | 5.49 | 101 | 5.2 | 75.92 |
| THP1_CTB_HIA_24_2 | 3.3 | 101 | 3.27 | 86.14 |
| THP1_CTB_HIA_24_3 | 2.93 | 101 | 2.89 | 86.32 |
| THP1_Con_4_1 | 3.32 | 101 | 3.2 | 78.84 |
| THP1_Con_4_2 | 1.5 | 101 | 1.44 | 76.73 |
| THP1_Con_4_3 | 3.95 | 101 | 3.82 | 79.53 |
| THP1_Con_24_1 | 6.2 | 101 | 5.92 | 77.71 |
| THP1_Con_24_2 | 3.06 | 101 | 2.94 | 81.42 |
| THP1_Con_24_3 | 4.11 | 101 | 3.98 | 80.57 |

**Table S2.** Read quality assessment statistics from FastQC analysis.

| Sample | Duplicates % | GC Content % | Average Sequence Length | Total Sequences | Fails % |
|---|---|---|---|---|---|
| HCjE_CTA_4_1_R 1 | 55.31 | 46 | 77.75 | 5437118 | 18.18 |
| HCjE_CTA_4_2_R 1 | 51.83 | 47 | 72.92 | 3666532 | 18.18 |
| HCjE_CTA_4_3_R 1 | 50.88 | 47 | 74.46 | 3678624 | 18.18 |
| HCjE_CTA_24_1_ R1 | 55.43 | 46 | 78.97 | 6695509 | 18.18 |
| HCjE_CTA_24_2_ R1 | 54.38 | 46 | 79.55 | 4472025 | 18.18 |
| HCjE_CTA_24_3_ R1 | 50.18 | 45 | 81.03 | 3995901 | 18.18 |
| HCjE_CTA_HIA_4 _1_R1 | 49.18 | 45 | 81.76 | 5010155 | 9.09 |
| HCjE_CTA_HIA_4 _2_R1 | 47.59 | 46 | 82.2 | 3249517 | 9.09 |
| HCjE_CTA_HIA_4 _3_R1 | 45.37 | 48 | 67.84 | 1163449 | 18.18 |
| HCjE_CTA_HIA_2 4_1_R1 | 54.19 | 46 | 78.14 | 5052717 | 18.18 |
| HCjE_CTA_HIA_2 4_2_R1 | 50.61 | 46 | 78.41 | 4000742 | 18.18 |
| HCjE_CTA_HIA_24_3_R1 | 48.15 | 44 | 76.63 | 4994200 | 9.09 |
| HCjE_CTB_4_1_R 1 | 48.53 | 45 | 80.96 | 4977697 | 9.09 |
| HCjE_CTB_4_2_R 1 | 53.1 | 46 | 79.31 | 4728873 | 18.18 |
| HCjE_CTB_4_3_R 1 | 49.93 | 46 | 77.77 | 3133953 | 9.09 |
| HCjE_CTB_24_1_ R1 | 53.33 | 47 | 77.82 | 4555754 | 18.18 |
| HCjE_CTB_24_2_ R1 | 48.85 | 44 | 81.92 | 4698958 | 9.09 |
| HCjE_CTB_24_3_ R1 | 53.09 | 48 | 76.26 | 4396168 | 27.27 |
| HCjE_CTB_HIA_4 _1_R1 | 53.73 | 46 | 79.08 | 5865167 | 18.18 |
| HCjE_CTB_HIA_4 _2_R1 | 50.18 | 46 | 78.69 | 3475796 | 18.18 |
| HCjE_CTB_HIA_4 _3_R1 | 49.12 | 46 | 80.37 | 3837505 | 9.09 |
| HCjE_CTB_HIA_2 4_1_R1 | 51.42 | 46 | 78.11 | 4131317 | 18.18 |
| HCjE_CTB_HIA_2 4_2_R1 | 55.73 | 47 | 76.51 | 4047313 | 27.27 |
| HCjE_CTB_HIA_2 4_3_R1 | 56.06 | 46 | 79.59 | 6193701 | 18.18 |
| HCjE_Con_4_1_R 1 | 55.05 | 45 | 81.17 | 6862080 | 18.18 |
| HCjE_Con_4_2_R 1 | 52.34 | 44 | 82.44 | 6216293 | 18.18 |
| HCjE_Con_4_3_R 1 | 47.76 | 45 | 80.57 | 3501732 | 9.09 |
| HCjE_Con_24_1_ R1 | 52.06 | 47 | 77.08 | 4268720 | 18.18 |
| HCjE_Con_24_2_ R1 | 49.44 | 46 | 75.85 | 3268176 | 9.09 |
| HCjE_Con_24_3_ R1 | 50.8 | 48 | 72.5 | 3259502 | 27.27 |
| THP1_CTA_4_1_R 1 | 56.33 | 46 | 78.59 | 4318690 | 27.27 |
| THP1_CTA_4_2_R 1 | 58.15 | 47 | 81.12 | 3771956 | 27.27 |
| THP1_CTA_4_3_R 1 | 59.16 | 47 | 82.49 | 3895121 | 27.27 |
| THP1_CTA_24_1_ R1 | 50.52 | 45 | 80.66 | 5389217 | 18.18 |
| THP1_CTA_24_2_ R1 | 48.85 | 45 | 80.16 | 3842924 | 9.09 |
| THP1_CTA_24_3_ R1 | 52.47 | 47 | 74.09 | 3258491 | 18.18 |
| THP1_CTA_HIA_4 _1_R1 | 56.14 | 46 | 82.6 | 3354842 | 27.27 |
| THP1_CTA_HIA_4 _2_R1 | 54.89 | 46 | 78.71 | 2866114 | 27.27 |
| THP1_CTA_HIA_4 _3_R1 | 52.48 | 45 | 76.18 | 4552992 | 18.18 |
| THP1_CTA_HIA_2 4_1_R1 | 49.75 | 47 | 74.98 | 2978089 | 9.09 |

| | | | | | |
|---|---|---|---|---|---|
| THP1_CTA_HIA_2 4_2_R1 | 50.21 | 46 | 77.73 | 3523239 | 18.18 |
| THP1_CTA_HIA_2 4_3_R1 | 57.15 | 46 | 80.31 | 5979177 | 18.18 |
| THP1_CTB_4_1_R 1 | 48.49 | 45 | 83.05 | 3879594 | 9.09 |
| THP1_CTB_4_2_R 1 | 48.26 | 45 | 78.09 | 4429426 | 9.09 |
| THP1_CTB_4_3_R 1 | 49.05 | 45 | 81.05 | 4214464 | 9.09 |
| THP1_CTB_24_1_ R1 | 48.09 | 46 | 80.93 | 3359761 | 9.09 |
| THP1_CTB_24_2_ R1 | 46.56 | 45 | 80.85 | 3198384 | 9.09 |
| THP1_CTB_24_3_ R1 | 51.37 | 45 | 78.65 | 4400766 | 18.18 |
| THP1_CTB_HIA_4 _1_R1 | 50.21 | 45 | 80.93 | 4703467 | 18.18 |
| THP1_CTB_HIA_4 _2_R1 | 50.75 | 47 | 73.92 | 3771046 | 18.18 |
| THP1_CTB_HIA_4 _3_R1 | 48.61 | 45 | 80.04 | 3933650 | 9.09 |
| THP1_CTB_HIA_2 4_1_R1 | 49.79 | 44 | 75.25 | 5203742 | 9.09 |
| THP1_CTB_HIA_2 4_2_R1 | 50.74 | 46 | 85.39 | 3266180 | 18.18 |
| THP1_CTB_HIA_2 4_3_R1 | 49.04 | 46 | 85.56 | 2889971 | 9.09 |
| THP1_Con_4_1_R 1 | 53.84 | 47 | 78.14 | 3202841 | 36.36 |
| THP1_Con_4_2_R 1 | 53.74 | 48 | 76.05 | 1443922 | 36.36 |
| THP1_Con_4_3_R 1 | 55.92 | 47 | 78.82 | 3817368 | 27.27 |
| THP1_Con_24_1_ R1 | 50.38 | 44 | 77.02 | 5924676 | 18.18 |
| THP1_Con_24_2_ R1 | 54.75 | 46 | 80.7 | 2944487 | 27.27 |
| THP1_Con_24_3_ R1 | 59.62 | 47 | 79.85 | 3978514 | 27.27 |

**Table S3.** Read alignment statistics from STAR mapping analysis.

| Sample | Number of input reads | Uniquely mapped reads number | Uniquely mapped reads % | Average input read length |
|---|---|---|---|---|
| HCjE_CTA_4_1 | 5437118 | 3459573 | 63.63 | 78 |
| HCjE_CTA_4_2 | 3666532 | 2342327 | 63.88 | 73 |
| HCjE_CTA_4_3 | 3678624 | 2355420 | 64.03 | 75 |
| HCjE_CTA_24_1 | 6695509 | 4334216 | 64.73 | 79 |
| HCjE_CTA_24_2 | 4472025 | 2907787 | 65.02 | 80 |
| HCjE_CTA_24_3 | 3995901 | 2727267 | 68.25 | 81 |
| HCjE_CTA_HIA_4 _1 | 5010155 | 3578217 | 71.42 | 82 |
| HCjE_CTA_HIA_4 _2 | 3249517 | 2244697 | 69.08 | 82 |
| HCjE_CTA_HIA_4 _3 | 1163449 | 732595 | 62.97 | 68 |
| HCjE_CTA_HIA_2 4_1 | 5052717 | 3358681 | 66.47 | 78 |
| HCjE_CTA_HIA_2 4_2 | 4000742 | 2663156 | 66.57 | 79 |
| HCjE_CTA_HIA_2 4_3 | 4994200 | 3446040 | 69 | 77 |
| HCjE_CTB_4_1 | 4977697 | 3495059 | 70.21 | 81 |
| HCjE_CTB_4_2 | 4728873 | 3041336 | 64.31 | 80 |
| HCjE_CTB_4_3 | 3133953 | 2033053 | 64.87 | 78 |
| HCjE_CTB_24_1 | 4555754 | 2916144 | 64.01 | 78 |
| HCjE_CTB_24_2 | 4698958 | 3306386 | 70.36 | 82 |
| HCjE_CTB_24_3 | 4396168 | 2664342 | 60.61 | 76 |
| HCjE_CTB_HIA_4 _1 | 5865167 | 3904464 | 66.57 | 79 |
| HCjE_CTB_HIA_4 _2 | 3475796 | 2243706 | 64.55 | 79 |
| HCjE_CTB_HIA_4 _3 | 3837505 | 2600818 | 67.77 | 81 |
| HCjE_CTB_HIA_2 4_1 | 4131317 | 2639202 | 63.88 | 78 |
| HCjE_CTB_HIA_2 4_2 | 4047313 | 2291698 | 56.62 | 77 |
| HCjE_CTB_HIA_2 4_3 | 6193701 | 4017067 | 64.86 | 80 |
| HCjE_Con_4_1 | 6862080 | 4690485 | 68.35 | 81 |
| HCjE_Con_4_2 | 6216293 | 4413019 | 70.99 | 83 |
| HCjE_Con_4_3 | 3501732 | 2436901 | 69.59 | 81 |
| HCjE_Con_24_1 | 4268720 | 2756849 | 64.58 | 77 |
| HCjE_Con_24_2 | 3268176 | 2050458 | 62.74 | 76 |
| HCjE_Con_24_3 | 3259502 | 1971042 | 60.47 | 73 |
| THP1_CTA_4_1 | 4318690 | 2684838 | 62.17 | 79 |
| THP1_CTA_4_2 | 3771956 | 2331887 | 61.82 | 81 |
| THP1_CTA_4_3 | 3895121 | 2444804 | 62.77 | 83 |
| THP1_CTA_24_1 | 5389217 | 4003579 | 74.29 | 81 |
| THP1_CTA_24_2 | 3842924 | 2870687 | 74.7 | 80 |
| THP1_CTA_24_3 | 3258491 | 2208119 | 67.77 | 74 |
| THP1_CTA_HIA_ 4_1 | 3354842 | 2168926 | 64.65 | 83 |
| THP1_CTA_HIA_ 4_2 | 2866114 | 1812289 | 63.23 | 79 |
| THP1_CTA_HIA_ 4_3 | 4552992 | 3081279 | 67.68 | 76 |
| THP1_CTA_HIA_ 24_1 | 2978089 | 2092390 | 70.26 | 75 |

| | | | | |
|---|---|---|---|---|
| THP1_CTA_HIA_ 24_2 | 3523239 | 2499126 | 70.93 | 78 |
| THP1_CTA_HIA_ 24_3 | 5979177 | 4237924 | 70.88 | 81 |
| THP1_CTB_4_1 | 3879594 | 2936903 | 75.7 | 83 |
| THP1_CTB_4_2 | 4429426 | 3333823 | 75.27 | 78 |
| THP1_CTB_4_3 | 4214464 | 3106703 | 73.72 | 81 |
| THP1_CTB_24_1 | 3359761 | 2385612 | 71.01 | 81 |
| THP1_CTB_24_2 | 3198384 | 2372941 | 74.19 | 81 |
| THP1_CTB_24_3 | 4400766 | 3137682 | 71.3 | 79 |
| THP1_CTB_HIA_4 _1 | 4703467 | 3520298 | 74.84 | 81 |
| THP1_CTB_HIA_4 _2 | 3771046 | 2567110 | 68.07 | 74 |
| THP1_CTB_HIA_4 _3 | 3933650 | 2872960 | 73.04 | 80 |
| THP1_CTB_HIA_2 4_1 | 5203742 | 3873484 | 74.44 | 75 |
| THP1_CTB_HIA_2 4_2 | 3266180 | 2387054 | 73.08 | 86 |
| THP1_CTB_HIA_2 4_3 | 2889971 | 2109283 | 72.99 | 86 |
| THP1_Con_4_1 | 3202841 | 1956446 | 61.08 | 78 |
| THP1_Con_4_2 | 1443922 | 831328 | 57.57 | 76 |
| THP1_Con_4_3 | 3817368 | 2385711 | 62.5 | 79 |
| THP1_Con_24_1 | 5924676 | 4330419 | 73.09 | 77 |
| THP1_Con_24_2 | 2944487 | 1955595 | 66.42 | 81 |
| THP1_Con_24_3 | 3978514 | 2449835 | 61.58 | 80 |

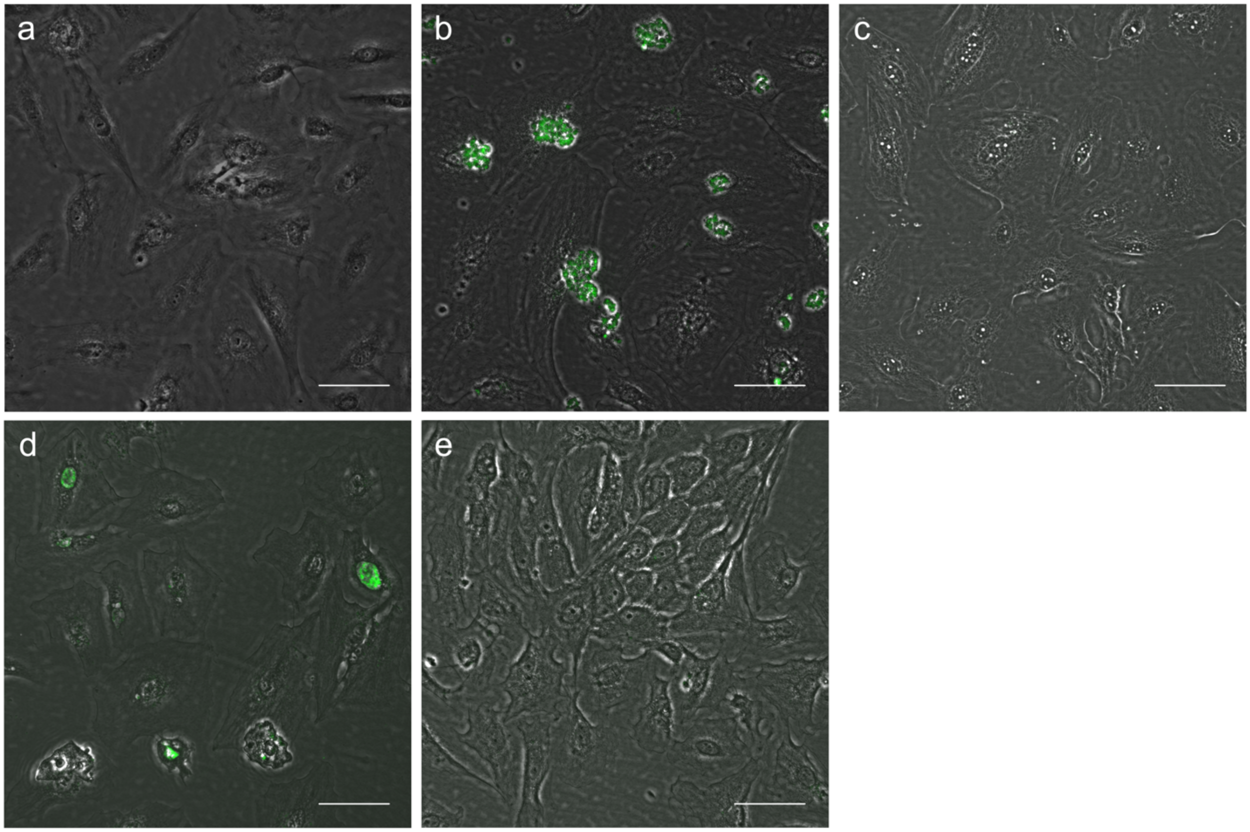


**Fig. S1.** Immunofluorescence detection of CtA and CtB in HCjE cells. Ct inclusions were visualised using anti-LPS antibodies (Pathfinder Chlamydia Confirmation System). (**a**) Uninoculated control, (**b**) live CtA, (**c**) HIA CtA, (d) live CtB, (**e**) HIA CtB. Scale bars = 50 µm.

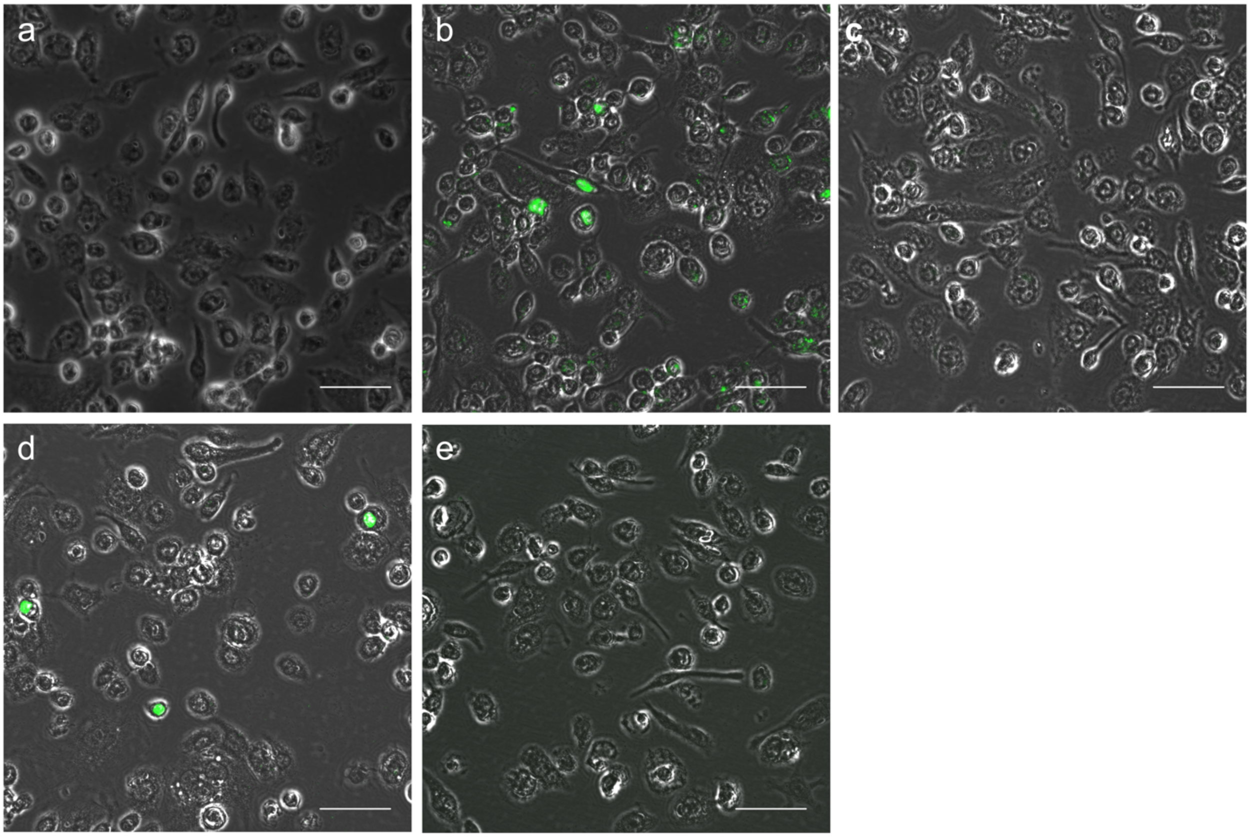


**Fig. S2.** Immunofluorescence detection of CtA and CtB in THP1 cells. Ct inclusions were visualised using anti-LPS antibodies (Pathfinder Chlamydia Confirmation System). (**a**) Uninoculated control, (**b**) live CtA, (**c**) HIA CtA, (d) live CtB, (**e**) HIA CtB. Scale bars = 50 µm.

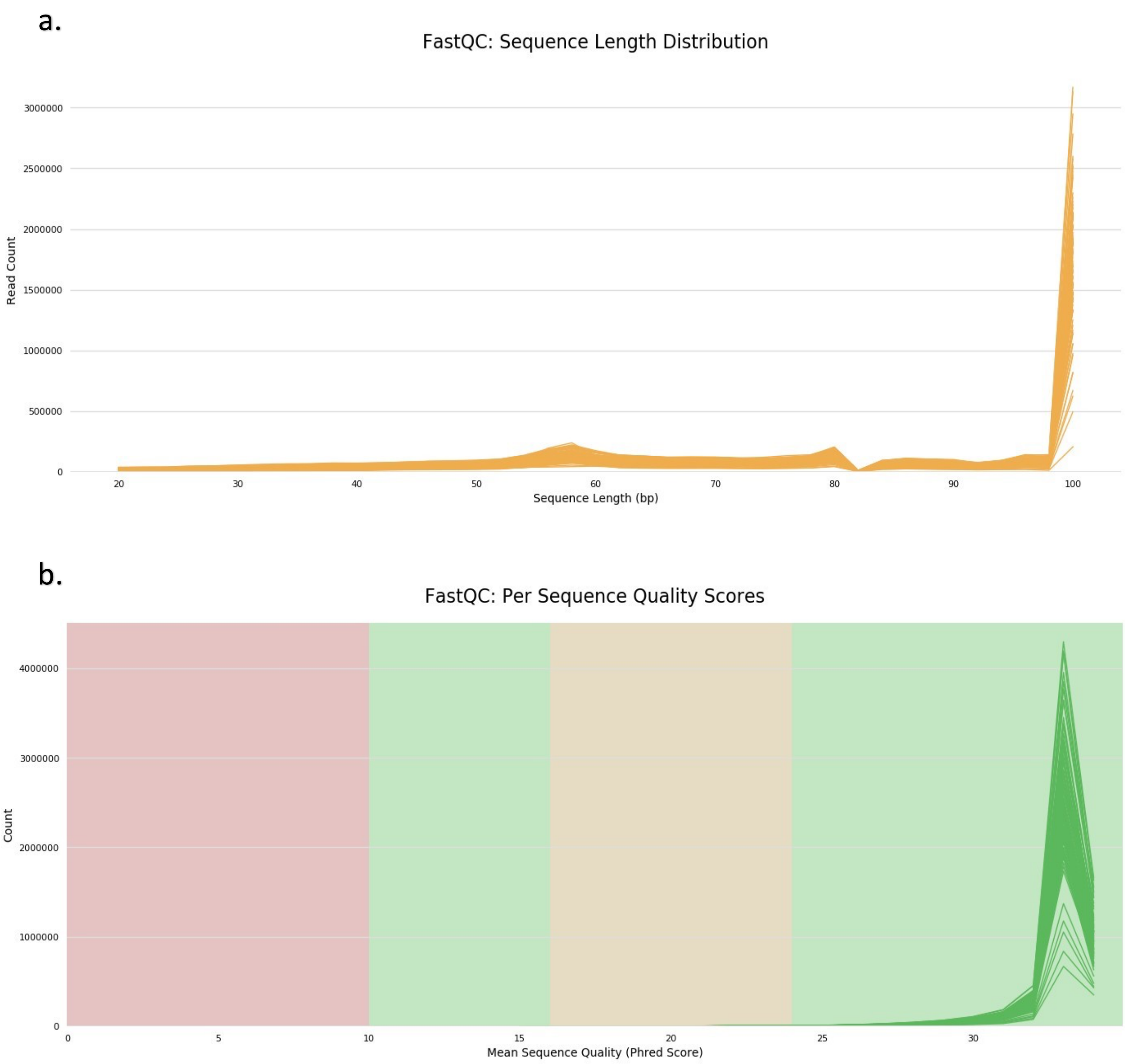


**Fig. S3.** (a) Read length distribution from FastQC analysis. (b) Per-read quality score distribution from FastQC analysis.

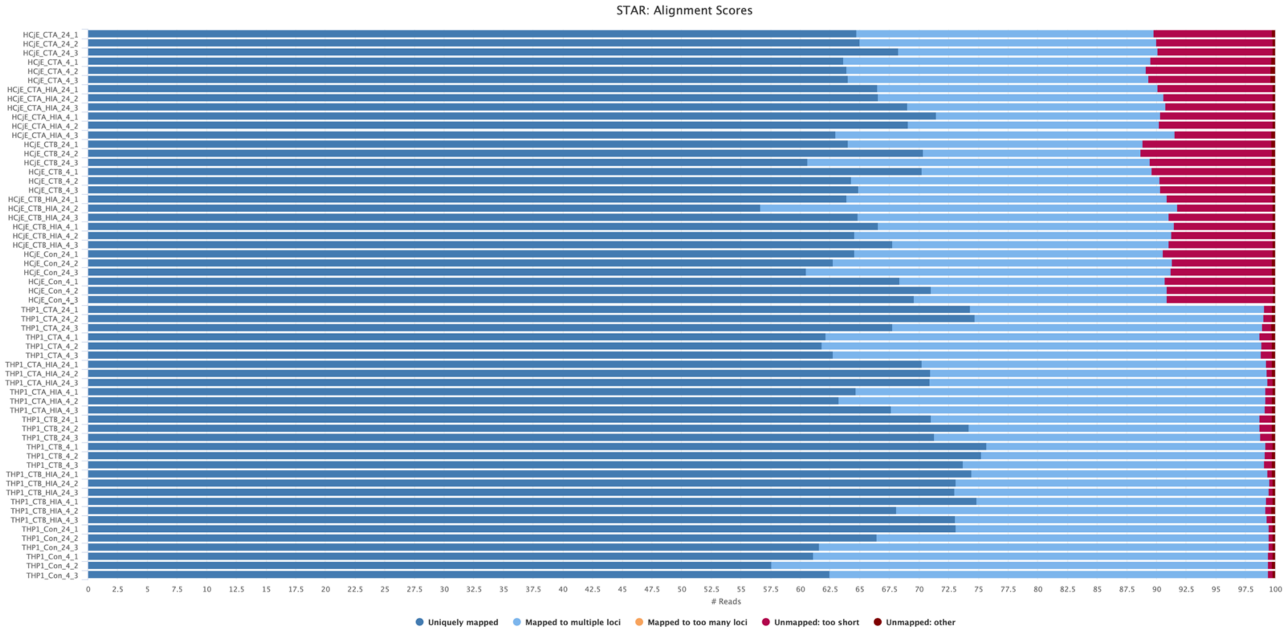


**Fig. S4.** Read alignment summary showing the proportion of reads in each alignment category per sample.

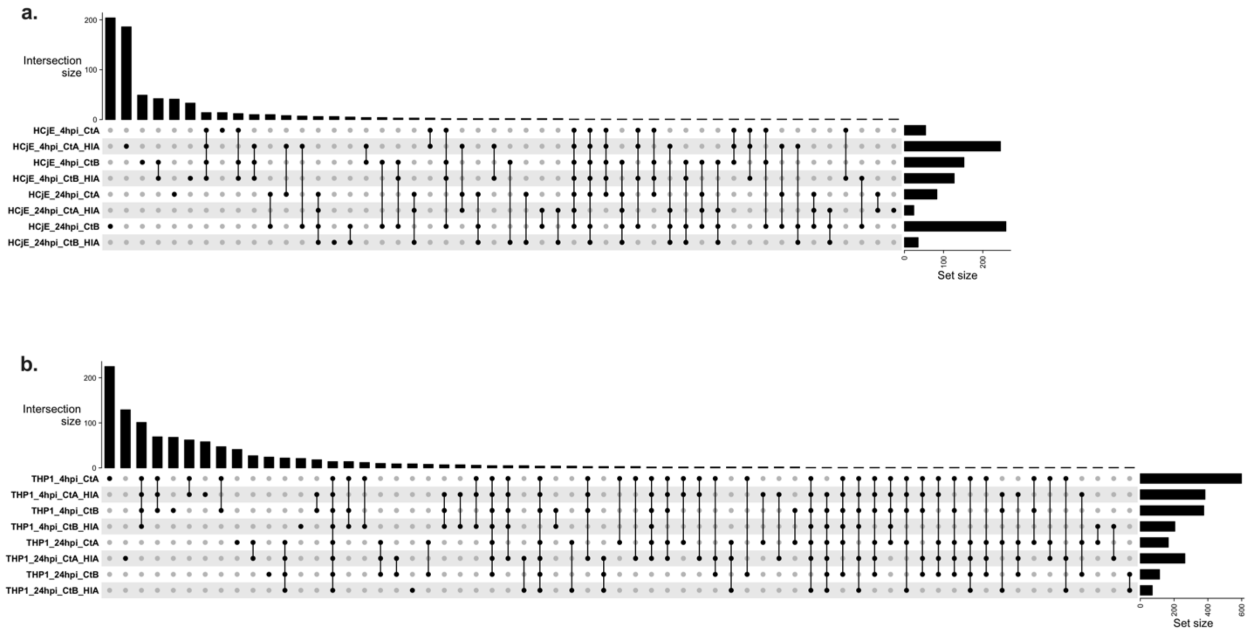


**Fig. S5.** UpSet plot showing the intersection of DEGs across various inoculation conditions in HCjE (**a**) and THP1 (**b**) cells. The plot visualises the number of DEGs unique to each condition and shared among combinations of conditions. Vertical bars represent the size of each intersection set, while filled dots below indicate which conditions are involved in each intersection.

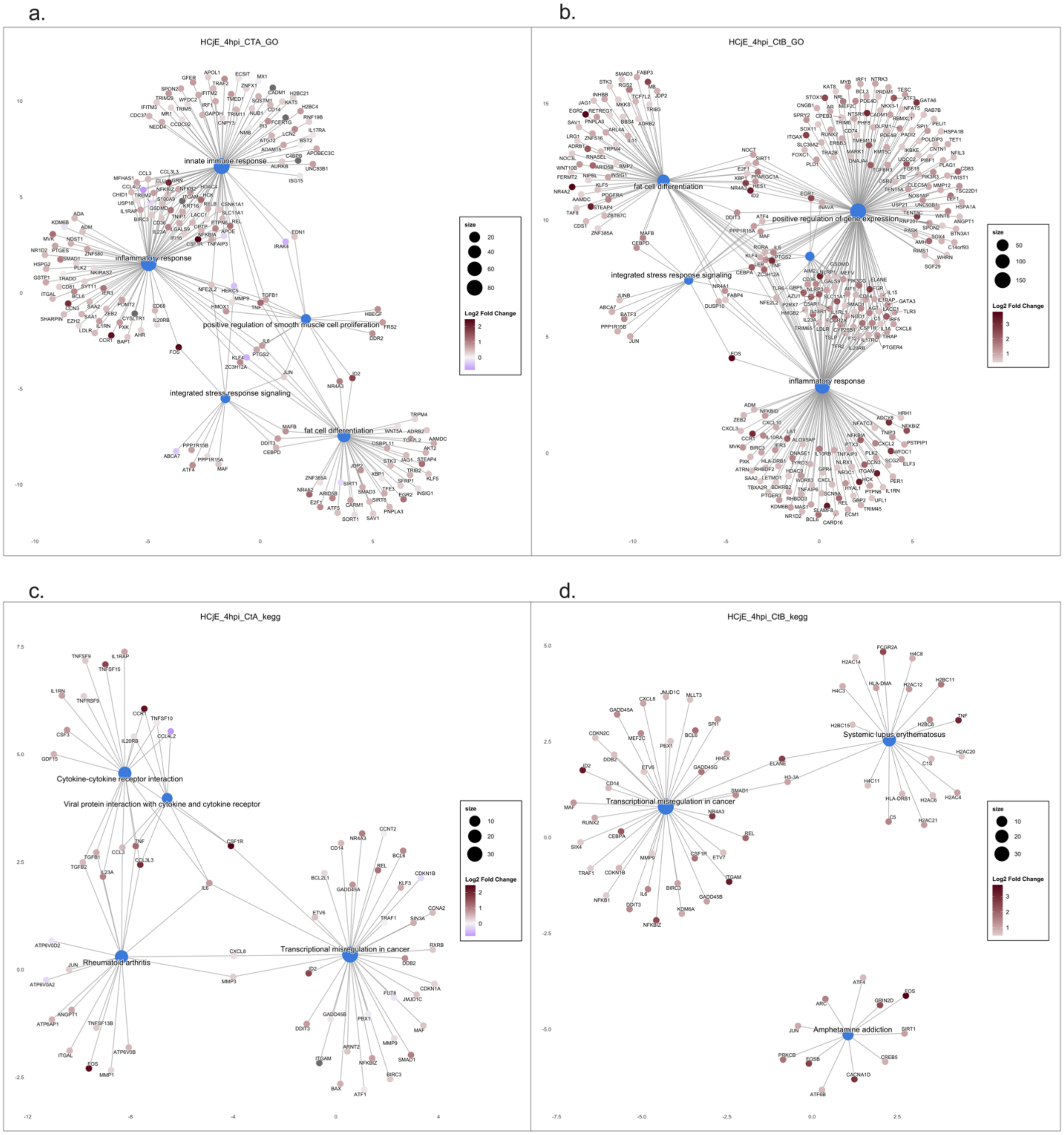
a.
HCjE_4hpi_CTA_GO
innate immune response
inflammatory response
positive regulation of smooth muscle cell proliferation
integrated stress response signaling
fat cell differentiation
b.
HCjE_4hpi_CtB_GO
fat cell differentiation
positive regulation of gene expression
integrated stress response signaling
inflammatory response
c.
HCjE_4hpi_CtA_kegg
Cytokine-cytokine receptor interaction
Viral protein interaction with cytokine and cytokine receptor
Rheumatoid arthritis
Transcriptional misregulation in cancer
d.
HCjE_4hpi_CtB_kegg
Transcriptional misregulation in cancer
Systemic lupus erythematosus
Amphetamine addiction
size
Log2 Fold Change

**Fig. S6.**

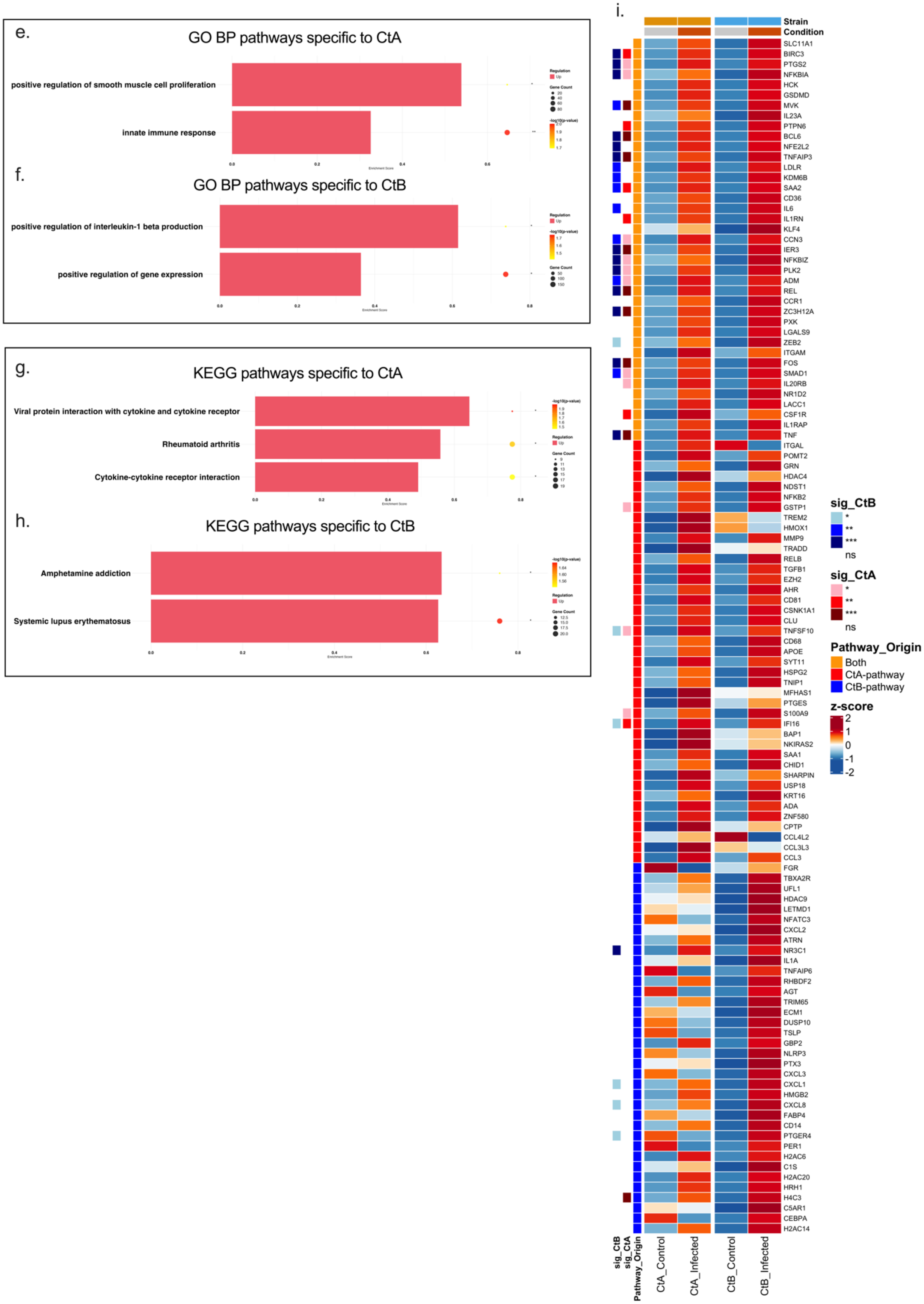
e.
GO BP pathways specific to CtA
positive regulation of smooth muscle cell proliferation
innate immune response
f.
GO BP pathways specific to CtB
positive regulation of interleukin-1 beta production
positive regulation of gene expression
g.
KEGG pathways specific to CtA
Viral protein interaction with cytokine and cytokine receptor
Rheumatoid arthritis
Cytokine-cytokine receptor interaction
h.
KEGG pathways specific to CtB
Amphetamine addiction
Systemic lupus erythematosus
i.
Strain
Condition
SLC11A1
BIRC3
PTGS2
NFKBIA
HCK
GSDMD
MVK
IL23A
PTPN6
BCL6
NFE2L2
TNFAIP3
LDLR
KDM6B
SAA2
CD36
IL6
IL1RN
KLF4
CCN3
IER3
NFKBIZ
PLK2
ADM
REL
CCR1
ZC3H12A
PXK
LGALS9
ZEB2
ITGAM
FOS
SMAD1
IL20RB
NR1D2
LACC1
CSF1R
IL1RAP
TNF
ITGAL
POMT2
GRN
HDAC4
NDST1
NFKB2
GSTP1
TREM2
HMOX1
MMP9
TRADD
RELB
TGFB1
EZH2
AHR
CD81
CSNK1A1
CLU
TNFSF10
CD68
APOE
SYT11
HSPG2
TNIP1
MFHAS1
PTGES
S100A9
IFI16
BAP1
NKIRAS2
SAA1
CHID1
SHARPIN
USP18
KRT16
ADA
ZNF580
CPTP
CCL4L2
CCL3L3
CCL3
FGR
TBXA2R
UFL1
HDAC9
LETMD1
NFATC3
CXCL2
ATRN
NR3C1
IL1A
TNFAIP6
RHBDF2
AGT
TRIM65
ECM1
DUSP10
TSLP
GBP2
NLRP3
PTX3
CXCL3
CXCL1
HMGB2
CXCL8
FABP4
CD14
PTGER4
PER1
H2AC6
C1S
H2AC20
HRH1
H4C3
C5AR1
CEBPA
H2AC14
sig_CtB
sig_CtA
Pathway_Origin
CtA_Control
CtA_Infected
CtB_Control
CtB_Infected
sig_CtB
*
**
***
ns
sig_CtA
*
**
***
ns
Pathway_Origin
Both
CtA-pathway
CtB-pathway
z-score
2
1
0
-1
-2

129

**Fig. S6.** Pathway enrichment analysis and differential gene expression profiles in HCjE cells infected with Ct strains A/2497 and B/Tunis864 at 4 hpi. (**a-d**) Pathway-gene interaction networks for significantly enriched GO BP (**a**, **b**) and KEGG (**c**, **d**) pathways following CtA and CtB infection. Pathway nodes are sized by significance; gene nodes are coloured by log2FC (blue: downregulated; red: upregulated). (**e-h**) Strain-specific pathway enrichment profiles for GO BP (**e**, **f**) and KEGG (**g**, **h**). Bars show enrichment magnitude/direction; overlaid circles represent gene count (size) and significance (colour intensity, -log10 adjusted *P*-value). (**i**) Heatmap of differentially expressed pathway-associated genes (adjusted P < 0.05). Columns show mean Z-scored expression for controls and infected samples (CtA: A/2497; CtB: B/Tunis864). Row annotations indicate pathway origin (red: CtA-pathway; blue: CtB-pathway; orange: both) and strain-specific significance levels. Column annotations denote condition (grey: control; orange: infected) and strain (orange: CtA; blue: CtB). Colour scale: blue (low) to red (high) expression. Significance: *** $P < 0.001$, ** $P < 0.01$, * $P < 0.05$.

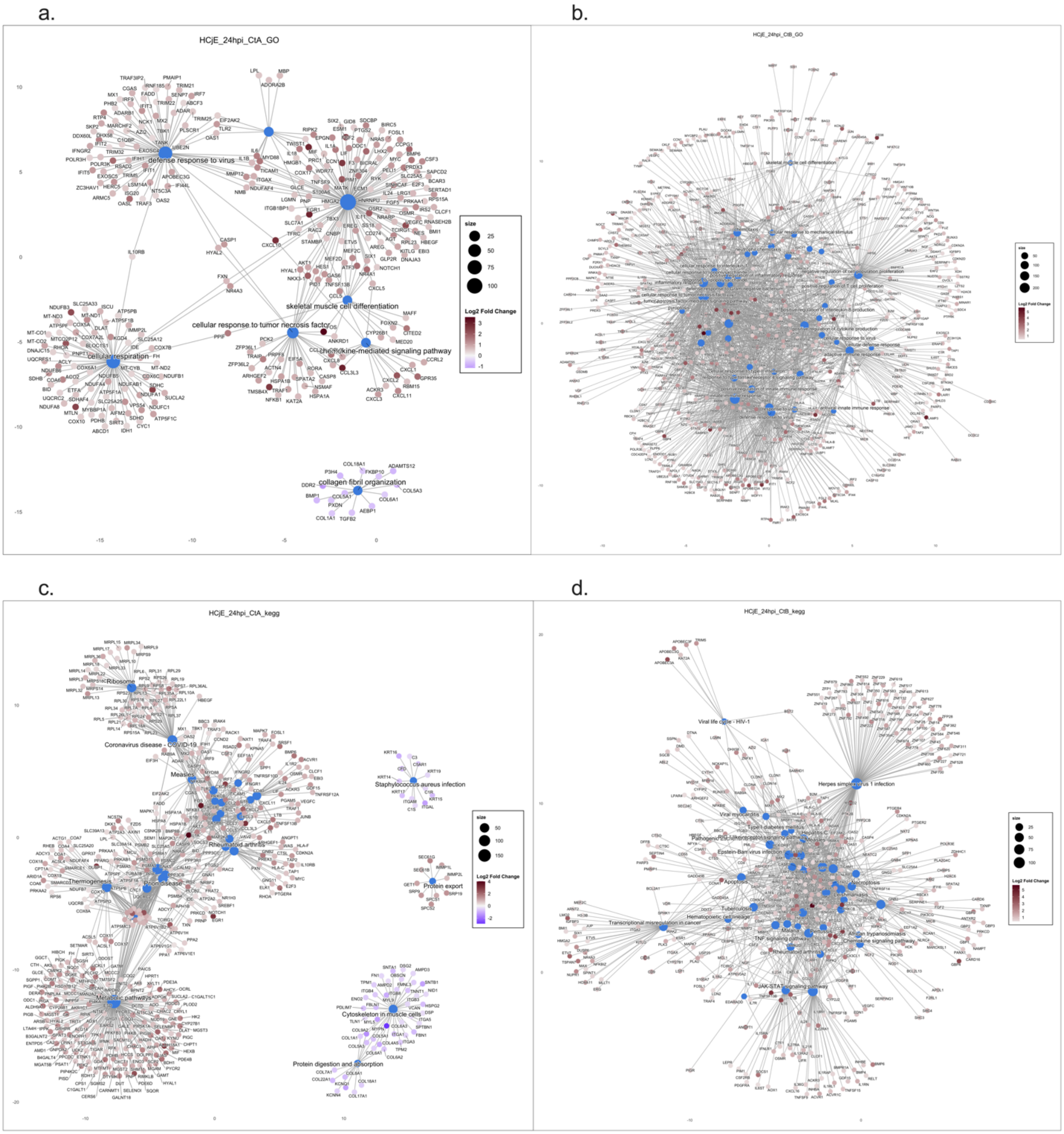
a.
HCjE_24hpi_CtA_GO
defense response to virus
cellular respiration
cellular response to tumor necrosis factor
skeletal muscle cell differentiation
chemokine-mediated signaling pathway
collagen fibril organization
size
Log2 Fold Change
b.
HCjE_24hpi_CtB_GO
c.
HCjE_24hpi_CtA_kegg
Ribosome
Coronavirus disease - COVID-19
Measles
Rheumatoid arthritis
Thermogenesis
Metabolic pathways
Staphylococcus aureus infection
Protein export
Cytoskeleton in muscle cells
Protein digestion and absorption
size
Log2 Fold Change
d.
HCjE_24hpi_CtB_kegg
Viral life cycle - HIV-1
Herpes simplex virus 1 infection
Epstein-Barr virus infection
Apoptosis
Tuberculosis
Transcriptional misregulation in cancer
Hematopoietic cell lineage
TNF signaling pathway
Malaria
African trypanosomiasis
Chemokine signaling pathway
JAK-STAT signaling pathway
size
Log2 Fold Change

**Fig. S7.**

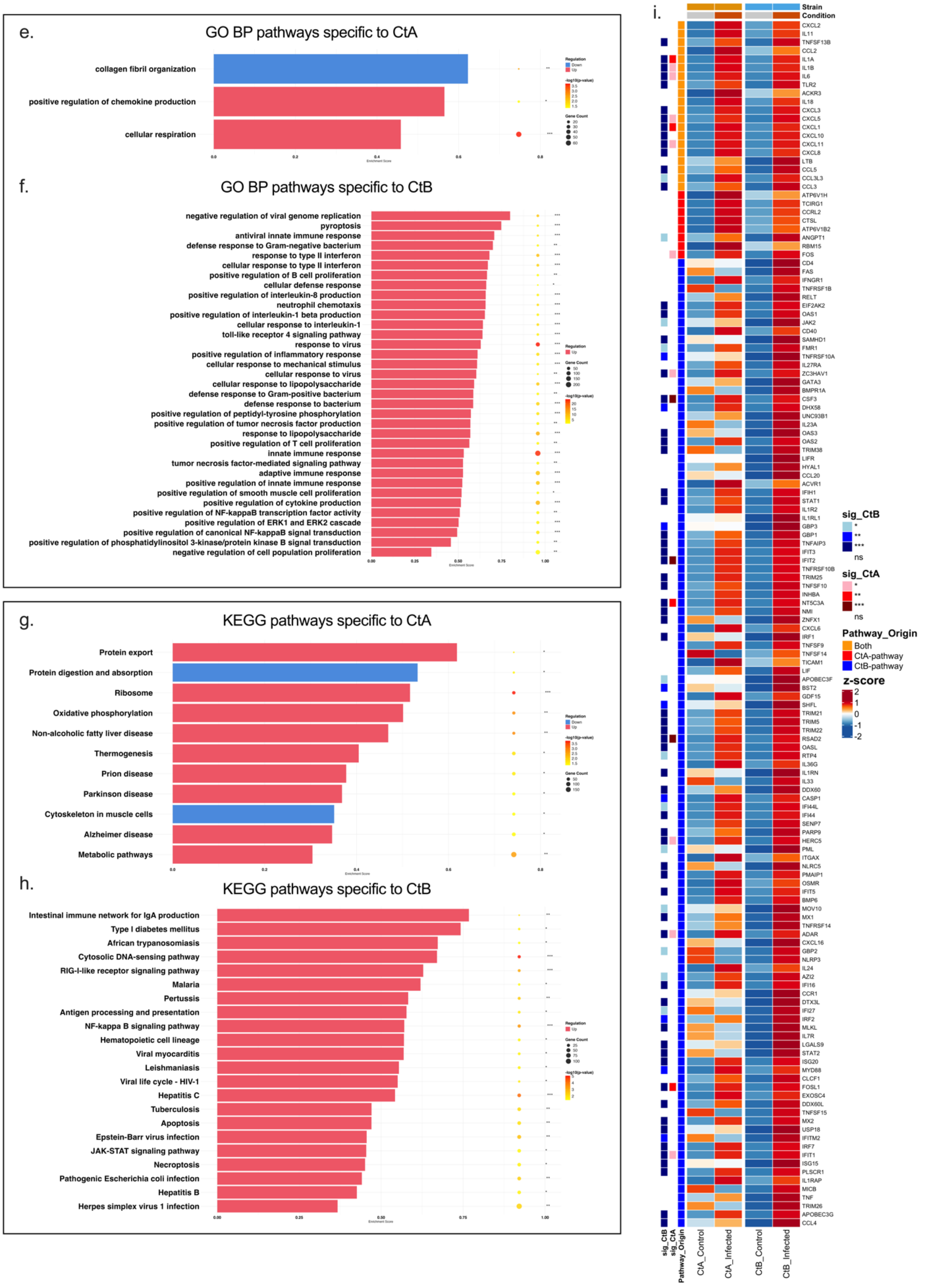
e.
GO BP pathways specific to CtA
collagen fibril organization
positive regulation of chemokine production
cellular respiration
f.
GO BP pathways specific to CtB
negative regulation of viral genome replication
pyroptosis
antiviral innate immune response
defense response to Gram-negative bacterium
response to type II interferon
cellular response to type II interferon
positive regulation of B cell proliferation
cellular defense response
positive regulation of interleukin-8 production
neutrophil chemotaxis
positive regulation of interleukin-1 beta production
cellular response to interleukin-1
toll-like receptor 4 signaling pathway
response to virus
positive regulation of inflammatory response
cellular response to mechanical stimulus
cellular response to virus
cellular response to lipopolysaccharide
defense response to Gram-positive bacterium
defense response to bacterium
positive regulation of peptidyl-tyrosine phosphorylation
positive regulation of tumor necrosis factor production
response to lipopolysaccharide
positive regulation of T cell proliferation
innate immune response
tumor necrosis factor-mediated signaling pathway
adaptive immune response
positive regulation of innate immune response
positive regulation of smooth muscle cell proliferation
positive regulation of cytokine production
positive regulation of NF-kappaB transcription factor activity
positive regulation of ERK1 and ERK2 cascade
positive regulation of canonical NF-kappaB signal transduction
positive regulation of phosphatidylinositol 3-kinase/protein kinase B signal transduction
negative regulation of cell population proliferation
g.
KEGG pathways specific to CtA
Protein export
Protein digestion and absorption
Ribosome
Oxidative phosphorylation
Non-alcoholic fatty liver disease
Thermogenesis
Prion disease
Parkinson disease
Cytoskeleton in muscle cells
Alzheimer disease
Metabolic pathways
h.
KEGG pathways specific to CtB
Intestinal immune network for IgA production
Type I diabetes mellitus
African trypanosomiasis
Cytosolic DNA-sensing pathway
RIG-I-like receptor signaling pathway
Malaria
Pertussis
Antigen processing and presentation
NF-kappa B signaling pathway
Hematopoietic cell lineage
Viral myocarditis
Leishmaniasis
Viral life cycle - HIV-1
Hepatitis C
Tuberculosis
Apoptosis
Epstein-Barr virus infection
JAK-STAT signaling pathway
Necroptosis
Pathogenic Escherichia coli infection
Hepatitis B
Herpes simplex virus 1 infection
i.
Strain
Condition
sig_CtB
sig_CtA
Pathway_Origin
Both
CtA-pathway
CtB-pathway
z-score
CtA_Control
CtA_Infected
CtB_Control
CtB_Infected

**Fig. S7.** Pathway enrichment analysis and differential gene expression profiles in HCjE cells infected with Ct strains A/2497 and B/Tunis864 at 24 hpi. (**a-d**) Pathway-gene interaction networks for significantly enriched GO BP (**a**, **b**) and KEGG (**c**, **d**) pathways following CtA and CtB infection. Pathway nodes are sized by significance; gene nodes are coloured by log2FC (blue: downregulated; red: upregulated). (**e-h**) Strain-specific pathway enrichment profiles for GO BP (**e**, **f**) and KEGG (**g**, **h**). Bars show enrichment magnitude/direction; overlaid circles represent gene count (size) and significance (colour intensity, -log10 adjusted *P*-value). (**i**) Heatmap of differentially expressed pathway-associated genes (adjusted P < 0.05). Columns show mean Z-scored expression for controls and infected samples (CtA: A/2497; CtB: B/Tunis864). Row annotations indicate pathway origin (red: CtA-pathway; blue: CtB-pathway; orange: both) and strain-specific significance levels. Column annotations denote condition (grey: control; orange: infected) and strain (orange: CtA; blue: CtB). Colour scale: blue (low) to red (high) expression. Significance: *** $P < 0.001$, ** $P < 0.01$, * $P < 0.05$.

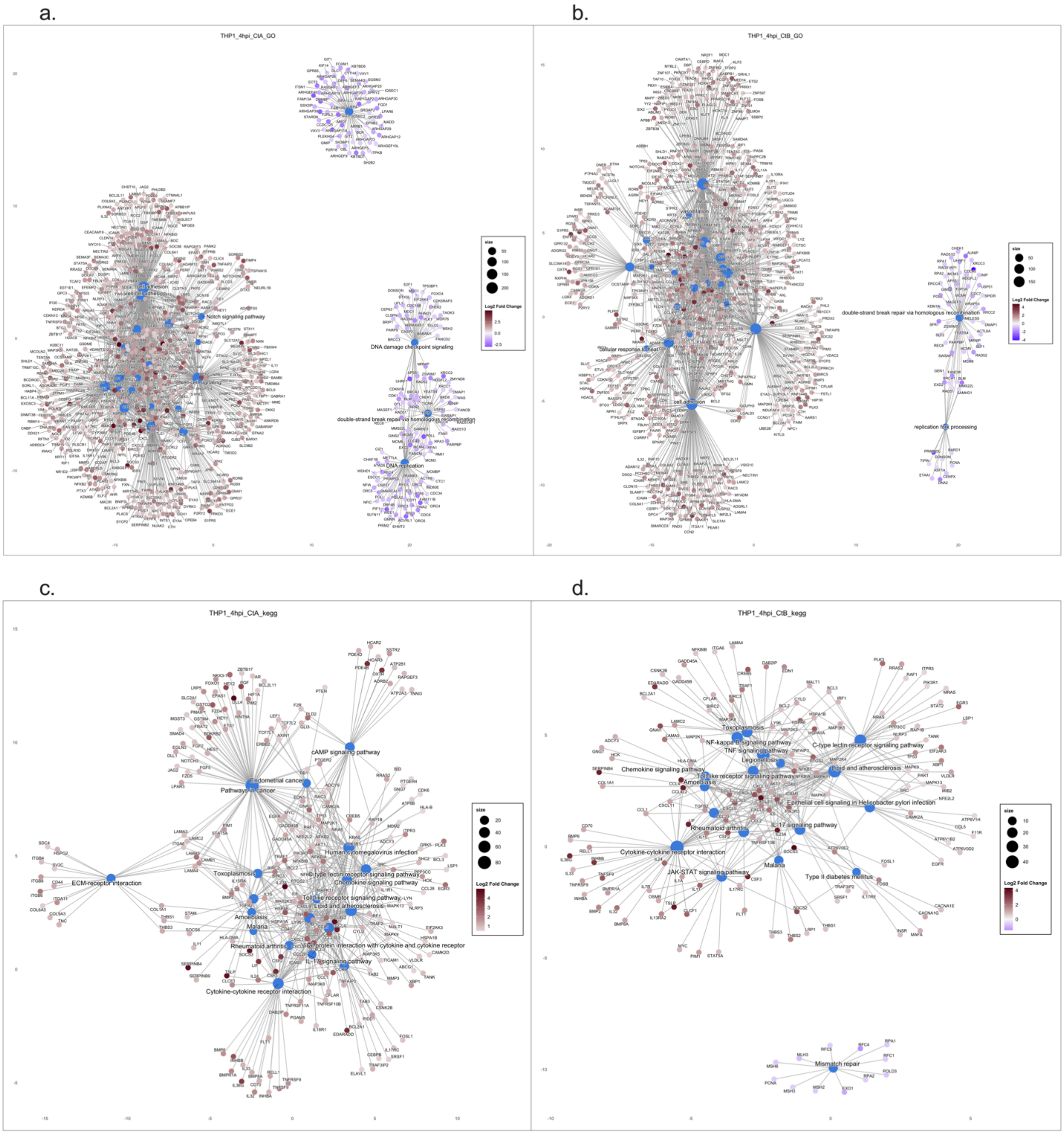
a.
THP1_4hpi_CtA_GO
Notch signaling pathway
DNA damage checkpoint signaling
double-strand break repair via homologous recombination
DNA replication
size
Log2 Fold Change
b.
THP1_4hpi_CtB_GO
cellular response to
double-strand break repair via homologous recombination
replication fork processing
size
Log2 Fold Change
c.
THP1_4hpi_CtA_kegg
cAMP signaling pathway
Endometrial cancer
Pathways in cancer
ECM-receptor interaction
Human cytomegalovirus infection
Toxoplasmosis
C-type lectin receptor signaling pathway
Chemokine signaling pathway
Toll-like receptor signaling pathway
Lipid and atherosclerosis
Amoebiasis
Malaria
Rheumatoid arthritis
IL-17 signaling pathway
Cytokine-cytokine receptor interaction
size
Log2 Fold Change
d.
THP1_4hpi_CtB_kegg
Toxoplasmosis
NF-kappa B signaling pathway
TNF signaling pathway
C-type lectin receptor signaling pathway
Legionellosis
Chemokine signaling pathway
Lipid and atherosclerosis
Amoebiasis
Toll-like receptor signaling pathway
Epithelial cell signaling in Helicobacter pylori infection
Rheumatoid arthritis
IL-17 signaling pathway
Cytokine-cytokine receptor interaction
JAK-STAT signaling pathway
Malaria
Type II diabetes mellitus
Mismatch repair
size
Log2 Fold Change

**Fig. S8.**

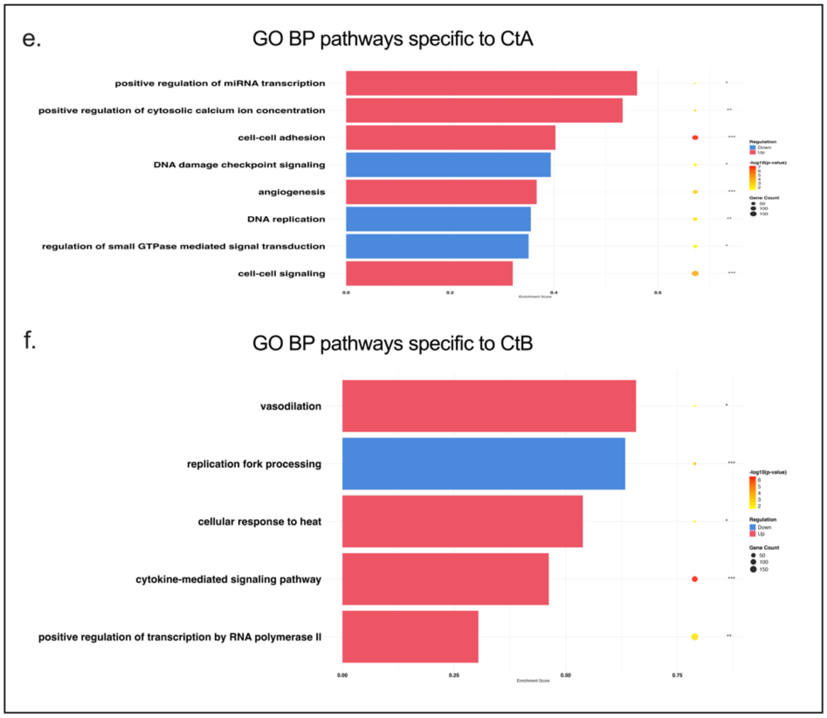
e.
GO BP pathways specific to CtA
positive regulation of miRNA transcription
positive regulation of cytosolic calcium ion concentration
cell-cell adhesion
DNA damage checkpoint signaling
angiogenesis
DNA replication
regulation of small GTPase mediated signal transduction
cell-cell signaling
f.
GO BP pathways specific to CtB
vasodilation
replication fork processing
cellular response to heat
cytokine-mediated signaling pathway
positive regulation of transcription by RNA polymerase II

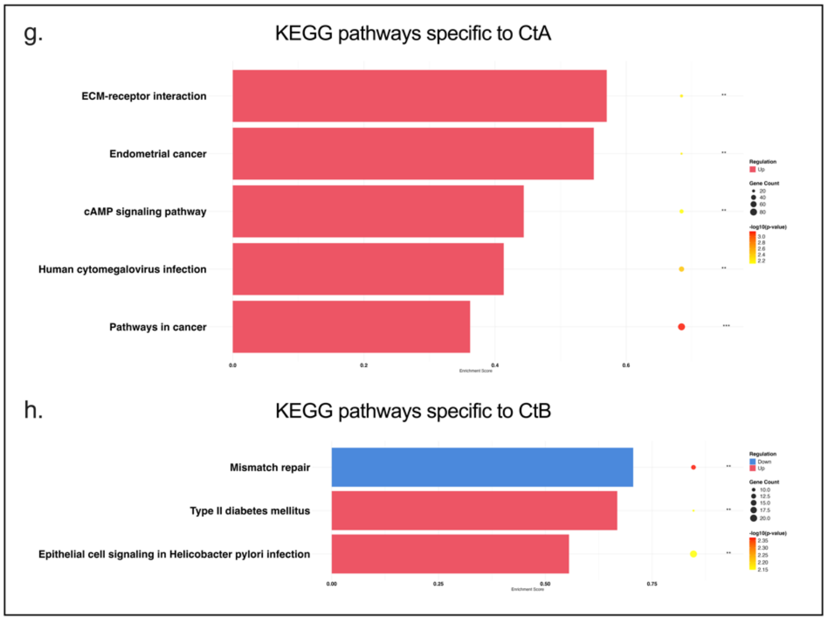
g.
KEGG pathways specific to CtA
ECM-receptor interaction
Endometrial cancer
cAMP signaling pathway
Human cytomegalovirus infection
Pathways in cancer
h.
KEGG pathways specific to CtB
Mismatch repair
Type II diabetes mellitus
Epithelial cell signaling in Helicobacter pylori infection

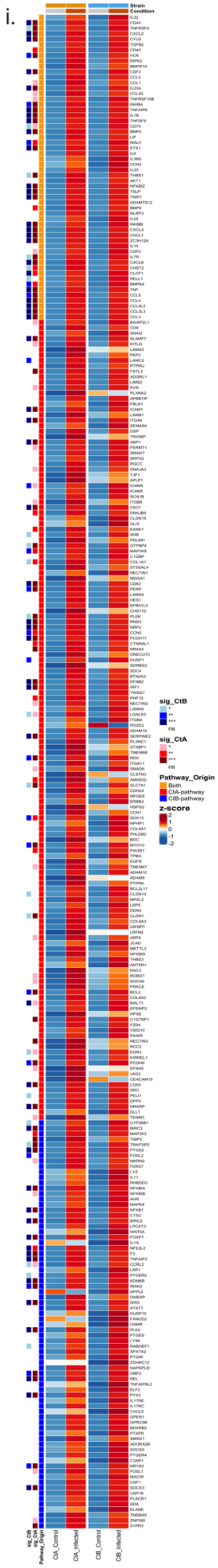
i.
sig_CtB
sig_CtA
Pathway_Origin
z-score

**Fig. S8.** Pathway enrichment analysis and differential gene expression profiles in THP-1 cells infected with Ct strains A/2497 and B/Tunis864 at 4 hpi. (**a-d**) Pathway-gene interaction networks for significantly enriched GO BP (**a**, **b**) and KEGG (**c**, **d**) pathways following CtA and CtB infection. Pathway nodes are sized by significance; gene nodes are coloured by log2FC (blue: downregulated; red: upregulated). (**e-h**) Strain-specific pathway enrichment profiles for GO BP (**e**, **f**) and KEGG (**g**, **h**). Bars show enrichment magnitude/direction; overlaid circles represent gene count (size) and significance (colour intensity, -log10 adjusted *P*-value). (**i**) Heatmap of differentially expressed pathway-associated genes (adjusted P < 0.05). Columns show mean Z-scored expression for controls and infected samples (CtA: A/2497; CtB: B/Tunis864). Row annotations indicate pathway origin (red: CtA-pathway; blue: CtB-pathway; orange: both) and strain-specific significance levels. Column annotations denote condition (grey: control; orange: infected) and strain (orange: CtA; blue: CtB). Colour scale: blue (low) to red (high) expression. Significance: *** *P* < 0.001, ** *P* < 0.01, * *P* < 0.05.

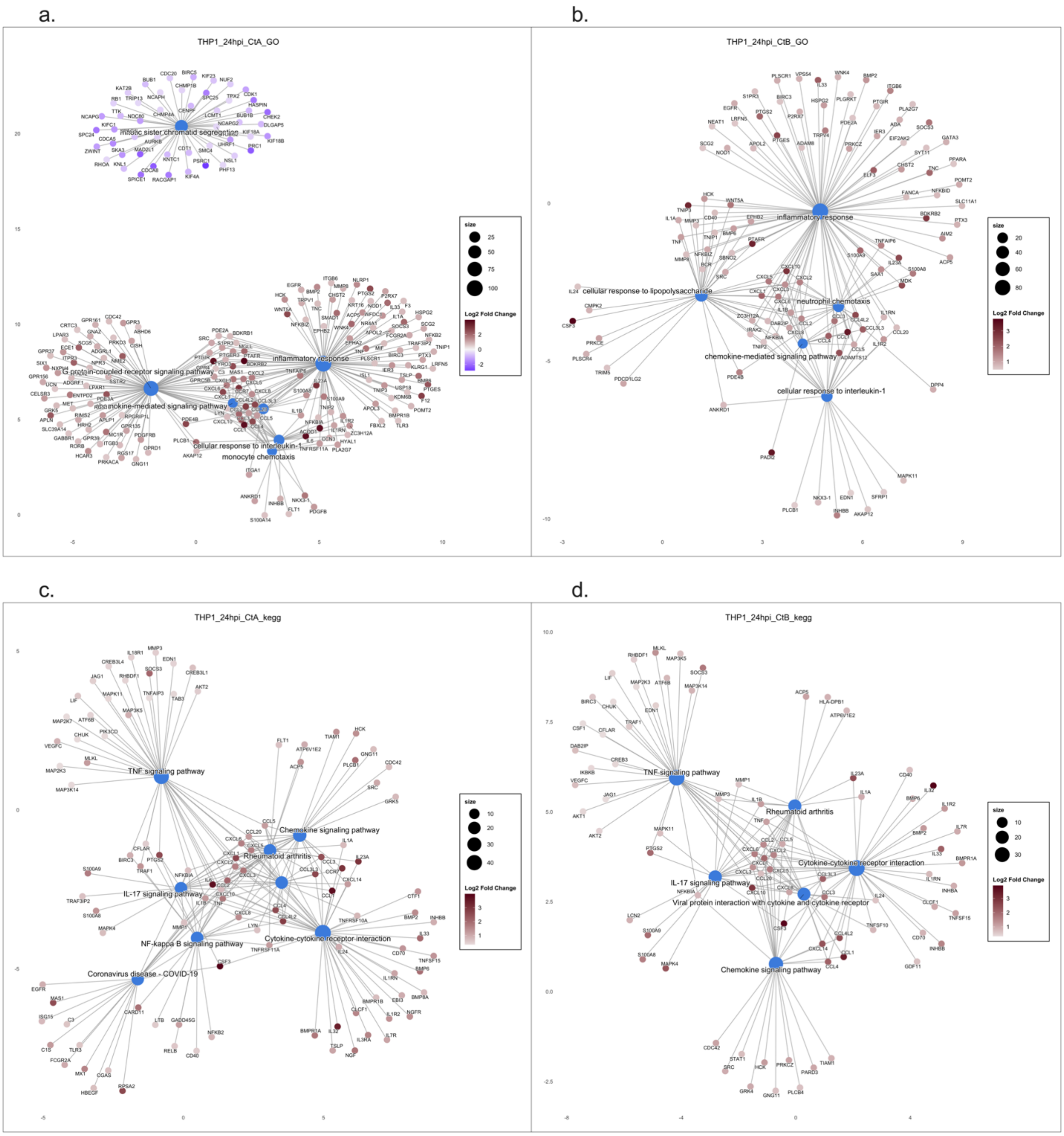
a.
THP1_24hpi_CtA_GO
mitotic sister chromatid segregation
G protein-coupled receptor signaling pathway
chemokine-mediated signaling pathway
inflammatory response
cellular response to interleukin-1
monocyte chemotaxis
size
Log2 Fold Change
b.
THP1_24hpi_CtB_GO
inflammatory response
cellular response to lipopolysaccharide
neutrophil chemotaxis
chemokine-mediated signaling pathway
cellular response to interleukin-1
size
Log2 Fold Change
c.
THP1_24hpi_CtA_kegg
TNF signaling pathway
Chemokine signaling pathway
Rheumatoid arthritis
IL-17 signaling pathway
NF-kappa B signaling pathway
Cytokine-cytokine receptor interaction
Coronavirus disease - COVID-19
size
Log2 Fold Change
d.
THP1_24hpi_CtB_kegg
TNF signaling pathway
Rheumatoid arthritis
Cytokine-cytokine receptor interaction
IL-17 signaling pathway
Viral protein interaction with cytokine and cytokine receptor
Chemokine signaling pathway
size
Log2 Fold Change

**Fig. S9.**

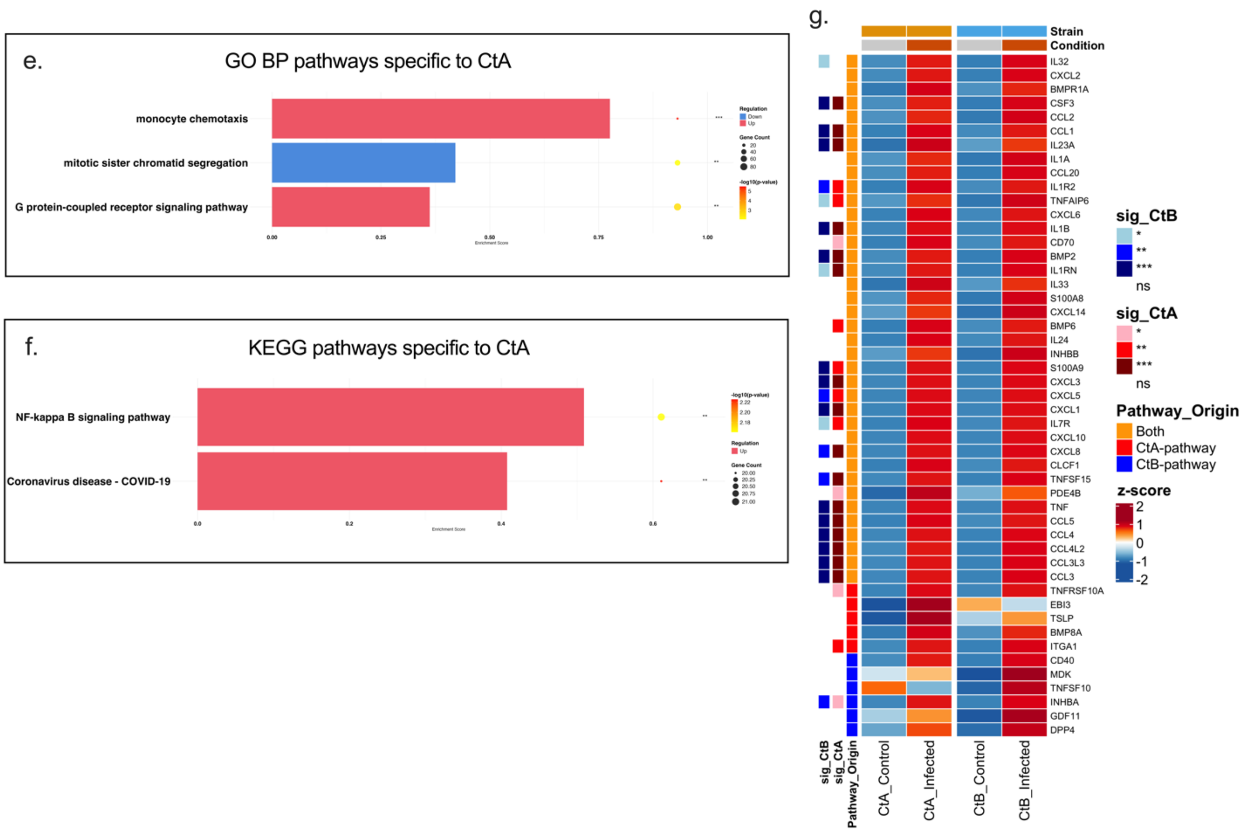


**Fig. S9.** Pathway enrichment analysis and differential gene expression profiles in THP-1 cells infected with Ct strains A/2497 and B/Tunis864 at 24 hpi. (**a-d**) Pathway-gene interaction networks for significantly enriched GO BP (**a**, **b**) and KEGG (**c**, **d**) pathways following CtA and CtB infection. Pathway nodes are sized by significance; gene nodes are coloured by log2FC (blue: downregulated; red: upregulated). (**e-f**) Strain-specific pathway enrichment profiles for GO BP (**e**) and KEGG (**f**). Bars show enrichment magnitude/direction; overlaid circles represent gene count (size) and significance (colour intensity, -log10 adjusted *P*-value). (**g**) Heatmap of differentially expressed pathway-associated genes (adjusted P < 0.05). Columns show mean Z-scored expression for controls and infected samples (CtA: A/2497; CtB: B/Tunis864). Row annotations indicate pathway origin (red: CtA-pathway; blue: CtB-pathway; orange: both) and strain-specific significance levels. Column annotations denote condition (grey: control; orange: infected) and strain (orange: CtA; blue: CtB). Colour scale: blue (low) to red (high) expression. Significance: *** $P < 0.001$, ** $P < 0.01$, * $P < 0.05$.
